# Higher baseline levels of fatty acid esters of hydroxy fatty acids do not further enhance the stimulatory effect of regular exercise on insulin sensitivity in obese mice

**DOI:** 10.64898/2026.06.08.730805

**Authors:** Marko Mitrovic, Olga Horakova, Martin Riecan, Veronika Kleinova, Petr Zouhar, Tomas Cajka, Ondrej Kuda, Lenka Rossmeislova, Martin Rossmeisl

## Abstract

**Background:** Exercise is an effective way to improve metabolic health, and the modulation of adipose tissue (AT) secretory functions may play a significant role in this process. AT produces various lipokines, including fatty acid esters of hydroxy fatty acids (FAHFA), which increase insulin sensitivity and have anti-inflammatory effects. While factors such as sex, age, obesity, and genetics influence FAHFA levels, their impact on exercise-induced FAHFA regulation remains unclear.

**Methods:** First, sex-specific responses to an acute bout of exercise were assessed in wild-type (WT) and ADTRP-deficient (ADTRP KO) mice. Fasted mice underwent acute treadmill exercise until exhaustion, followed by analysis of non-esterified fatty acids in plasma, *ex vivo* lipolysis in the presence or absence of a hormone-sensitive lipase (HSL) inhibitor, and FAHFA release from AT (measured by LC-MS). Second, obese male WT and ADTRP KO mice fed a high-fat diet underwent 7 weeks of regular treadmill exercise (5 days/week), after which parameters of glucose homeostasis, plasma and AT FAHFA levels, and AT lipid profiles were analyzed.

**Results:** Acute exercise-induced increases in plasma non-esterified fatty acid levels, AT lipolysis, and FAHFA release from AT explants were more pronounced in male mice of both genotypes. Conversely, pharmacological inhibition of HSL using BAY 59-9435 increased FAHFA release from AT explants only in females. In obese sedentary ADTRP KO mice, insulin sensitivity was improved compared with their WT counterparts. Although regular exercise suppressed weight gain in obese animals of both genotypes, insulin sensitivity improved only in WT mice. Chronic exercise generally had no effect on plasma FAHFA levels in mice fed *ad libitum*; however, in WT mice, it increased the levels of FAHFA-containing triacylglycerol estolides, which were associated with improved insulin sensitivity.

**Conclusions:** Acute exercise revealed sex-specific differences in AT lipolysis and FAHFA metabolism, with HSL playing an important role in FAHFA hydrolysis. Chronic exercise in obesity increases insulin sensitivity and FAHFA storage in AT; however, this effect is absent in ADTRP KO mice, which exhibit elevated FAHFA levels in AT, a condition associated with improved insulin sensitivity even in non-exercising animals.

## Background

Regular exercise is an important contributor to metabolic health, exerting beneficial effects on whole-body energy homeostasis and insulin sensitivity (1, 2). Some of these benefits are mediated through functional changes in adipose tissue (AT), a dynamic endocrine organ that regulates energy balance and systemic metabolism through a variety of mechanisms, including the secretion of bioactive substances such as lipokines (3–5). Both acute and chronic exercise influence AT function, with changes in lipokine secretion affecting downstream signaling pathways at both the tissue and systemic levels (6).

In 2014, a new class of lipokines called fatty acid esters of hydroxy fatty acids (FAHFA), predominantly synthesized in AT, was identified; the most extensively studied group within this class is palmitic acid esters of hydroxy stearic acids (PAHSA; (7)). These lipokines improve glucose homeostasis and insulin sensitivity (7), partly by suppressing lipolysis and hepatic glucose production (8), while also exerting anti-inflammatory effects in AT (7, 9). Other subclasses, such as linoleic acid- and docosahexaenoic acid-containing FAHFA, also display anti-inflammatory properties (10, 11). Importantly, not only various FAHFA species, but also different FAHFA regioisomers appear to have distinct biological effects; the impact of regioisomers on glucose homeostasis and inflammation depends on the position of the “branching” carbon (12). In AT, besides their free form, FAHFA are stored bound to glycerol in triacylglycerols, i.e., triacylglycerols estolides (TG EST; (13, 14)). Nevertheless, the specific biological functions of many FAHFA species have not yet been fully elucidated.

Studies indicate that FAHFA levels in AT are influenced by various biological and metabolic factors, such as sex, nutritional status, or genotype. For example, it has been found that female mice have higher concentrations of FAHFA in AT than males (15). In both subcutaneous and gonadal AT, fasting led to an increase in the levels of multiple FAHFA species (7, 14, 16, 17), and higher AT FAHFA levels were also observed in mice deficient in androgen-dependent TFPI-regulating protein (ADTRP), one of the enzymes involved in FAHFA degradation (15, 18, 19). Furthermore, PAHSA levels in humans were reduced in both AT and the bloodstream under pathophysiological conditions linked to insulin resistance and AT dysfunction (7, 20); FAHFA levels may also be influenced by diet and age (21). Regarding the effects of chronic exercise, we have previously demonstrated that 4 months of exercise training increased levels of most PAHSA regioisomers and their total levels in AT and the bloodstream in overweight older women (22). Furthermore, it has been shown that changes in blood FAHFA levels induced by acute exercise in trained runners allow for the differentiation between participants of normal weight and those who are overweight (23). However, it is not yet clear whether the release of FAHFA from AT triggered by acute exercise depends on sex, whether regular exercise can positively influence FAHFA metabolism in AT in cases of established obesity, and whether these effects are modulated in situations of persistently elevated FAHFA levels in AT, as is the case in mice lacking a functional ADTRP gene (i.e., in ADTRP KO mice).

Therefore, in this study, we first investigated the sex-specific effects of acute exercise on lipolysis and FAHFA release from AT in both wild-type (WT) and ADTRP KO mice, and subsequently analyzed the effect of a 7-week exercise training program on FAHFA metabolism in obese WT and ADTRP KO mice fed a high-fat diet.

## Materials and methods

### Animals and diets

A cohort of ADTRP KO mice and their WT littermates on a C57BL/6J genetic background (18, 19) was established at the Institute of Physiology of the Czech Academy of Sciences in Prague. All mice were housed individually under standard conditions, i.e., on a 12-hour light/dark cycle, at 22 °C, and with *ad libitum* access to water and food. Male and female WT and ADTRP KO mice were fed a standard diet (Rat/Mouse-Maintenance Extrudate; ssniff Spezialdiäten GmbH, Soest, Germany) until 4-5 months of age, when acute exercise experiments were conducted. For the chronic exercise experiments, male mice were fed a standard diet (see above) until 2 months of age, after which they were switched to a high-fat diet (HFD; “DIO-60 kJ% fat (Lard)”; ssniff Spezialdiäten GmbH, Soest, Germany).

### Experimental design and exercise protocol

A six-lane treadmill (Exer-3/6; Columbus Instruments, Columbus, OH, USA) was used for all exercise experiments. To investigate the potential influence of sex differences on the metabolic response to acute exercise, WT and ADTRP KO mice were food deprived from 11:00 AM to 3:00 PM and then subjected to the following acute exercise protocol: running at a speed of 8 m/min on a 5% incline for 40 min, followed by gradual increases of 0.5 m/min every 10 min until exhaustion (**Fig. 1A**). The exercising mice (EXE; *n* = 6) were killed and dissected immediately after completion of the protocol. Mice from the control group (CON; *n* = 7) remained in their cages without access to food or water for a period corresponding to the duration of the exercise session. To investigate the effects of chronic exercise, WT and ADTRP KO males were fed HFD for 6 weeks; thereafter, a 7-week intervention was initiated, consisting of regular exercise 5 days a week (Mon-Fri), while the mice continued to receive HFD (**Fig. 4A**). Every training day at 2:00 PM, the exercising mice (EXE, *n*=6) began running for 1 hour at a speed of 7 m/min on a 5% incline; the running speed was increased by 0.5 m/min each week and remained unchanged during the last two weeks of the intervention period (**Fig. 4B**). Sedentary controls (SED; *n*=7) were deprived of food and water for a period corresponding to the duration of daily exercise. An insulin tolerance test (ITT) was performed during the final week of the experimental protocol. Following the training period, mice rested for 3 days prior to the end of the experiment. Animal experiments were approved by the Institutional Animal Care and Use Committee and the Committee for Animal Protection of the Czech Academy of Sciences (Approval Number: 62/2023), in accordance with the EU Directive 2010/63/EU on the protection of animals used for scientific purposes.

**Figure 1.**
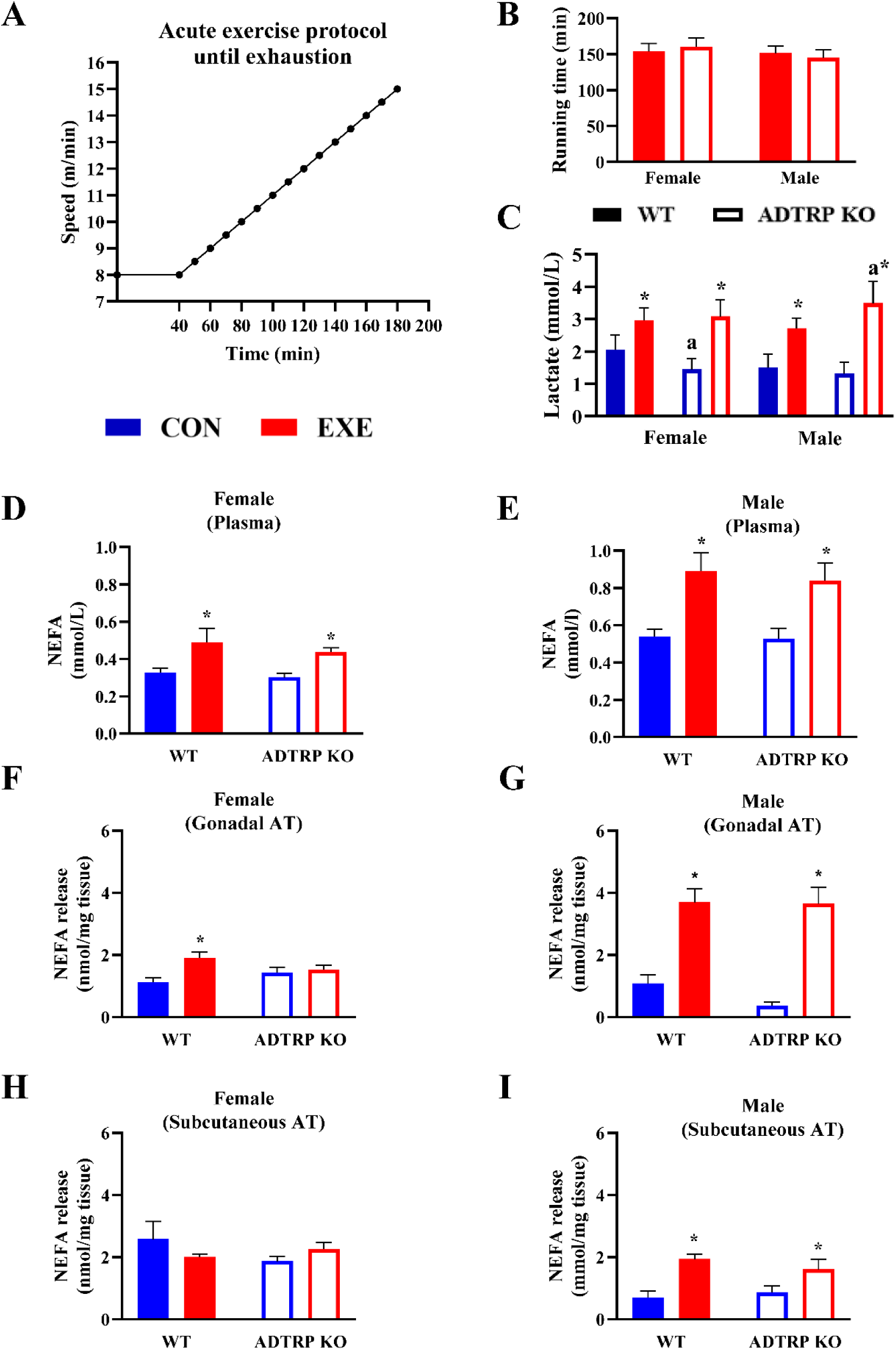
The effect of acute exercise on lipolysis-related parameters in male and female WT and ADTRP KO mice. A: Acute exercise protocol. **B:** Running time to exhaustion. **C:** Plasma lactate levels. **D-E:** Plasma NEFA levels in female (D) and male (E) WT and ADTRP KO mice. **F-I:** Release of NEFA from gonadal (F-G) and subcutaneous (H-I) AT explants of female (F and H) and male (G and I) mice. ^a,^*Significant difference vs. WT and CON, respectively; *p*≤0.05 (two-way ANOVA).

### Plasma and tissue collection

The mice were killed and dissected in the morning (9:00 AM) after having unrestricted access to food. Mice were killed by cervical dislocation under isoflurane anesthesia. Plasma was isolated after collecting the truncal blood into EDTA-containing tubes. Liver and AT samples were dissected and weighed. AT aliquots were immediately processed for explants, or snap-frozen in liquid N₂ and stored at -80 °C for later analyses. Samples designated for lipidomic analyses were kept on dry ice during the dissection and subsequently stored at -80 °C.

### Adipose tissue explants and ex vivo lipolysis

Aliquots (∼20 mg) of visceral (gonadal) or subcutaneous AT were washed with phosphate-buffered saline and preincubated on a 96-well plate in 200 μL of Dulbecco’s Modified Eagle’s Medium (DMEM) - low glucose (1 g/L; Gibco, UK), containing 2% fatty acid-free serum albumin (MP Biochemicals; Irvine, CA, USA), at 37°C in a 5% CO_2_ atmosphere for 1 hour. The explants (*n*=3 per mouse) were then transferred to new wells and incubated for another hour using the same medium and conditions as above. To inhibit hormone-sensitive lipase (HSL) activity, 10 μM BAY 59-9435 (MedChemExpress, Monmouth Junction, NJ, USA) was used, using a set of AT explants obtained from mice that had undergone acute exercise. The amount of non-esterified fatty acids (NEFA) released into the medium was measured using the NEFA-HR(2) kit from FUJIFILM Wako Chemicals Europe GmbH (Neuss, Germany).

### Circulating levels of metabolites and hormones

Blood glucose and lactate levels were measured in tail blood using the Contour Plus glucometers (Bayer Healthcare AG; Leverkusen, Germany) and the Lactate Plus Meter (Nova Biomedical; Waltham, MA, USA), respectively. Plasma levels of total cholesterol and total triacylglycerols were measured using commercially available colorimetric assays, Bio-La-Test TG L250S and Bio-La-Test CHOL L250S (Erba Lachema, Brno, Czech Republic), while plasma NEFA levels were determined using the kit from Wako Chemicals Europe GmbH (see above). Plasma insulin levels were measured using an Ultrasensitive Mouse Insulin ELISA kit (Mercodia; Uppsala, Sweden). The Homeostatic Model Assessment of Insulin Resistance (HOMA-IR) was applied to determine the degree of insulin resistance, using the following formula: fasting plasma insulin (mU/L)×fasting plasma glucose (mmol/L)/22.5. Adipose tissue IR (Adipo-IR) was calculated as fasting plasma NEFA × fasting plasma insulin measured after a 6-hour fast before the ITT (24).

### Body composition assessment

Total body fat mass, fat-free (lean) mass, and bone mineral content were assessed by dual-energy X-ray absorptiometry (DXA) using an InAlyzer densitometer (software version 3.2.3; MEDIKORS Inc., Gyeonggi, Korea). *In vivo* DXA measurements were performed during the final week of the chronic exercise program, with mice under light isoflurane anesthesia.

### Insulin tolerance test

The ITT was performed using intraperitoneal insulin injections (Actrapid Penfill, 100 IU/mL; Novo Nordisk, Bagsvaerd, Denmark). Insulin was administered at a dose of 0.4 U/kg body weight to fasted mice (from 6:00 AM to 12:00 PM).

### Tissue lipid content

The lipid content in homogenates of liver samples (aliquots of ∼50 mg) was determined by measuring tissue triacylglycerols, as before (25).

### Gene expression analysis

Gene expression in AT was analyzed using quantitative real-time PCR, as described previously (25). Transcript levels were normalized using the expression of the eukaryotic translation elongation factor 2 (*Eef2*) housekeeping gene. Primer sequences are listed in Supplementary Table 1 (**Table S1**).

### Untargeted lipidomic analysis and triacylglycerol estolide analysis

Lipidomic profiling of AT samples was performed using an untargeted liquid chromatography-mass spectrometry (LC-MS) workflow for the analysis of lipidome, as reported previously (26, 27). Details on the annotated lipid species are provided in **Table S5**.

### FAHFA regioisomer analysis

Samples of medium from AT explants (150 μL), plasma (150 μL), and AT (∼50 mg) were extracted according to previously described methods, with several modifications made to improve extraction yield and FAHFA detection during LC-MS analysis (7, 15). For details on the methodology, please refer to the Supplementary Information. The list of FAHFA species annotated in the media, plasma, and AT samples is provided in **Tables S2, S3** and **S4**, respectively.

### Data processing and statistics

Results are means ± SE. Differences were considered significant when *p* ≤ 0.05. Outliers were identified using Grubbs’ test and were excluded from subsequent analyses if statistically significant (*p* ≤ 0.05). Data normality was assessed using the Shapiro-Wilk test and through visual inspection of Q-Q plots. Variables that deviated from normality were subjected to log₁₀ transformation to better satisfy the assumptions underlying parametric statistical analyses. Two-way ANOVA followed by Fisher’s LSD post-hoc test was used to analyze the effect of genotype and exercise (GraphPad Prism 10.2. software; La Jolla, CA, USA). The Pearson correlation coefficient was used to assess the strength of the relationships between the selected variables. Lipidomic data were normalized using log_10_ transformation and Pareto scaling and analyzed by t-test performed in MetaboAnalyst 6.0 (28).

## Results

### Acute exercise caused a greater increase in adipose tissue lipolysis in male mice

The body composition of WT and ADTRP KO mice differed primarily by sex; females had significantly lower body weight and AT mass, regardless of genotype (**Tables S6 and S7**). This sexual dimorphism was reflected in lower circulating NEFA levels in female mice under basal conditions (**Fig. 1D,E**). All mice underwent an acute bout of moderate-intensity, prolonged (∼2.5 hours) exercise until exhaustion, with comparable running times across all groups (**Fig. 1B**) and similar post-exercise elevations in blood lactate (**Fig. 1C**) and glucose levels (**Tables S6 and S7**). Compared with sedentary controls, acute exercise also increased plasma NEFA levels in mice of both sexes and genotypes (**Fig. 1D,E**); however, pronounced sex- and genotype-dependent differences were observed in AT lipolysis, as determined by NEFA release from AT explants. In females, exercise increased NEFA release only in gonadal AT of WT animals (**Fig. 1F**), whereas no exercise-induced effects were observed in subcutaneous AT of mice of either genotype (**Fig. 1H**). In contrast, in males, acute exercise robustly stimulated the release of NEFA from both gonadal (**Fig. 1G**) and subcutaneous (**Fig. 1I**) AT, with comparable responses observed across both genotypes. In short, while male mice of both genotypes exhibit a robust exercise-induced lipolytic response, the loss of functional ADTRP prevents this response in females.

### Acute exercise led to increased FAHFA release from adipose tissue exclusively in males

We next evaluated the rate of FAHFA release from gonadal AT explants and identified the individual FAHFA molecules released (**Fig. 2** and **Fig. S1**). A total of 27 FAHFA regioisomers were detected in the conditioned medium. Of the various regioisomers detected in the medium under basal conditions, the most abundant in both females (**Fig. S1A**) and males (**Fig. S1B**) of both genotypes were members of the PAHSA family, particularly 9-PAHSA and 10-PAHSA. In females, the main factor influencing FAHFA secretion from the AT was genotype, not exercise (**Fig. 2A**). In contrast, in males, the total amount of FAHFA released from AT explants into the medium increased in response to acute exercise, with no differences observed between WT and ADTRP KO mice (**Fig. 2B**). Consistent with total FAHFA levels, the levels of individual FAHFA regioisomers in females did not change in response to acute exercise (except for 13-SAHLA, which decreased modestly in WT animals; **Fig. 2C**). In male mice, although exercise increased the total amount of FAHFA released from AT, this effect was driven by increases in a relatively limited subset of FAHFA regioisomers, particularly those enriched in oleic acid (OA) and/or linoleic acid (LA), i.e., 9-OAHOA, 11/10-LAHOA, 9-OAHLA, 13-OAHLA, 13-POHLA, 13-LAHLA (**Fig. 2D**). Thus, the profile of secreted FAHFA observed in gonadal AT explants from mice that underwent acute exercise did not fully overlap with the FAHFA profile that dominated the basal state in control, non-exercising mice (**Fig. 2** and **Fig. S1**). Overall, acute exercise regulates FAHFA release from AT in sex- and genotype-dependent manner.

**Figure 2.**
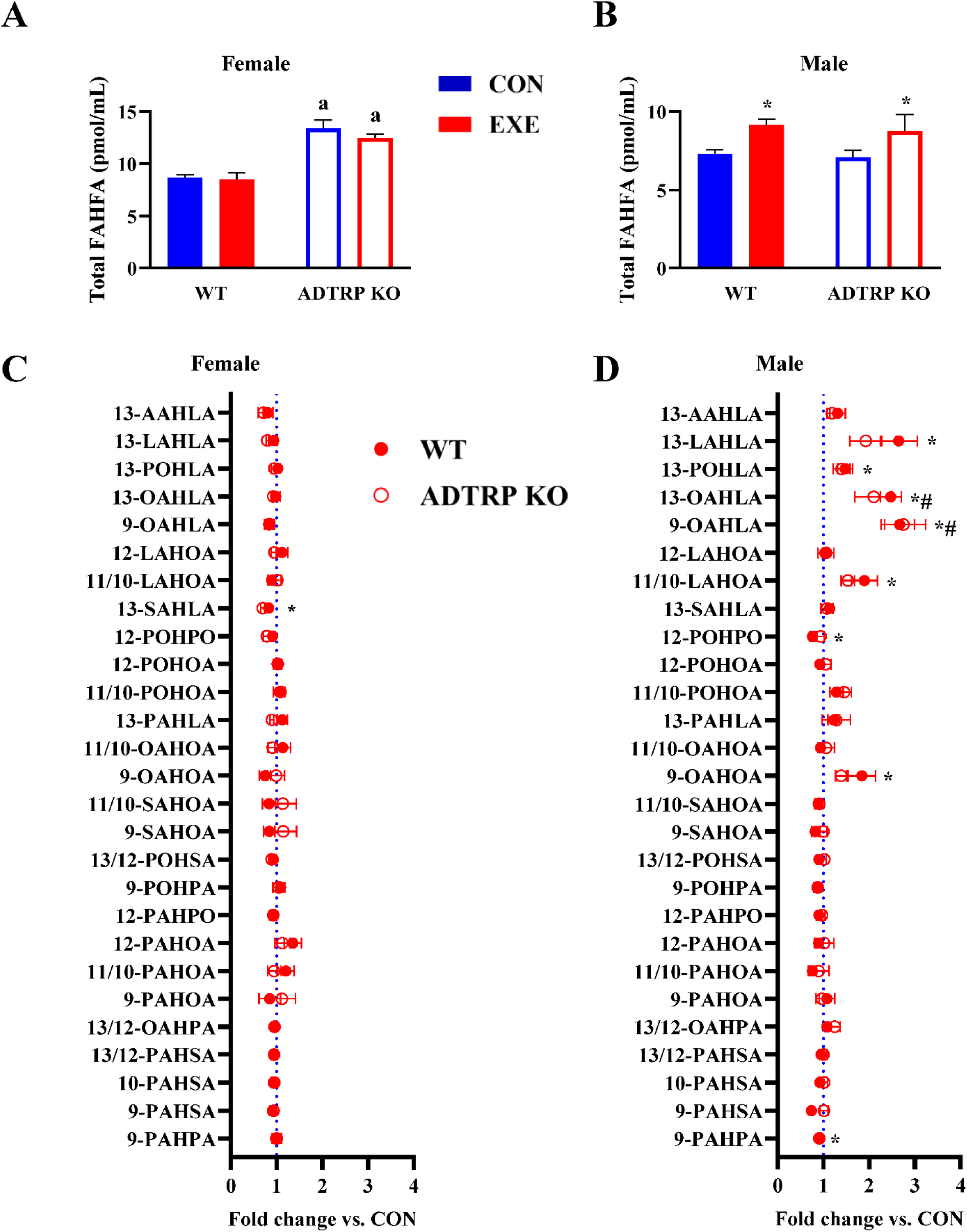
The sex-specific effects of acute exercise on FAHFA release from gonadal AT of WT and ADTRP KO mice. A-B: Total FAHFA released from AT explants from female (A) and male (B) mice subjected or not to a single bout of exercise. ^a,^*Significant difference vs. WT and CON, respectively; *p* ≤ 0.05 (two-way ANOVA). **C-D:** Levels of individual FAHFA regioisomers released from AT explants of female (C) and male (D) EXE mice of both genotypes (fold change vs. corresponding CON animals). *^,#^Significant difference vs. CON in WT and ADTRP KO group, respectively; *p*≤0.05 (*t*-test).

### HSL inhibition had opposite effects on FAHFA release from adipose tissue in male and female mice

Given that HSL has a dual role, i.e., it can release FAHFA from TG ESTs and catalyze their breakdown, we sought to determine which of these opposing functions predominates during acute exercise. To this end, we pharmacologically inhibited HSL in gonadal AT explants using the compound BAY 59-9435 and investigated how this inhibition affects the release of NEFA and FAHFA from gonadal AT explants of exercising mice of both sexes and genotypes. The HSL inhibition had no significant effect on NEFA release into the medium (**Fig. 3A,B**), except for a slight decrease observed in ADTRP KO females (**Fig. 3A**). In contrast, HSL inhibition revealed distinct sex-specific patterns in FAHFA mobilization following acute exercise (**Fig. 3C,D**). In WT females, HSL inhibition significantly increased total FAHFA release from AT explants (**Fig. 3C**); conversely, in WT males, HSL inhibition led to a significant decrease in FAHFA release, suggesting that in males, HSL primarily facilitated the mobilization of FAHFA from their TG EST (**Fig. 3D**). In AT explants from ADTRP KO mice, changes in FAHFA release following HSL inhibition were like those observed in their WT counterparts, but did not reach statistical significance (**Fig. 3C,D**). Regarding changes in the concentration of individual FAHFA species/regioisomers released into the medium because of HSL inhibition, most of them showed an increase in females compared to untreated controls, with statistically significant changes observed primarily in WT animals (**Fig. 3E**). In male-derived AT explants, HSL inhibition was associated with reduced levels of most secreted FAHFA regioisomers, particularly in WT mice (**Fig. 3F**). Notably, in males, 13-AAHLA was the only regioisomer for which this general downward trend was not observed (**Fig. 3F**). Overall, these results indicate that HSL function in FAHFA metabolism varies by sex: in females, this enzyme primarily regulates the breakdown of FAHFA, whereas in males, it facilitates their release from TG EST.

**Figure 3.**
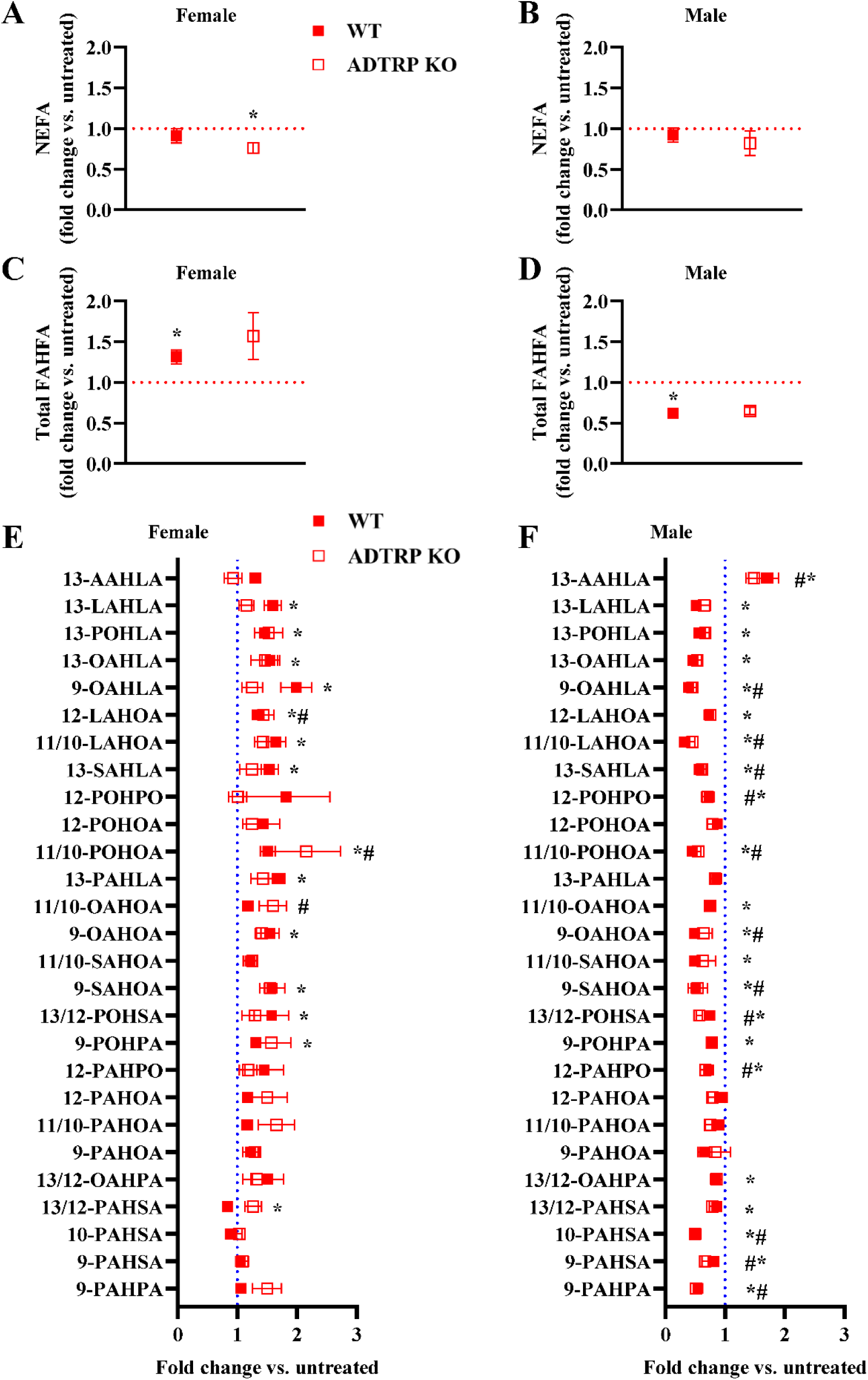
**Sex-specific effects of pharmacological inhibition of HSL on NEFA and FAHFA release from gonadal AT of exercising WT and ADTRP KO mice. A-B**: Release of NEFA from AT explants of female (A) and male (B) mice treated with the HSL inhibitor BAY 59-9435 (fold change vs. untreated explants from the same mice). *Significant difference vs. untreated explants; *p*≤0.05 (*t*-test). **C-D:** Total FAHFA release from AT explants of female (C) and male (D) mice treated with BAY 59-9435 (fold change vs. untreated explants). *Significant difference vs. untreated explants; *p*≤0.05 (*t*-test). **E-F:** Levels of individual FAHFA regioisomers released from AT explants of female (E) and male (F) mice treated with BAY 59-9435 (fold change vs. untreated explants). FAHFA species were sorted by total number of double bonds. *^,#^Significant difference vs. untreated explants in WT and ADTRP KO mice respectively; *p*≤0.05 (*t*-test).

### Chronic exercise suppressed further weight gain and improved glucose homeostasis in obese WT and ADTRP KO mice

Our next objective was to determine whether chronic exercise could improve AT function and FAHFA metabolism in mice with existing obesity induced by HFD feeding, also in situations where FAHFA levels in AT are constitutively elevated (i.e., in ADTRP KO mice). Given the significant contribution of HSL and its hydrolase activity towards FAHFA in the metabolic response to acute exercise in females (**Fig. 2** and **Fig. 3**), we used only male WT and ADTRP KO mice in this study. Regular exercise over a 7-week period suppressed weight gain in obese HFD-fed mice of both genotypes (**Table 1**), with differences becoming apparent as early as 3 weeks after initiation of the training regime (**Fig. 4C,D**). As expected, the EXE mice had lower body fat mass than the SED controls, as revealed by DEXA analysis (**Table 1**). In addition, the WT EXE mice also exhibited increased bone mineral content (**Table 1**). Food intake during the experiment was not affected by either genotype or exercise (**Fig. S2A**). Furthermore, significantly lower glucose levels were observed in EXE mice of both genotypes at all measured time points during the ITT (**Fig. 4E,F**). However, compared with the corresponding SED controls, the area under the curve (AUC) calculated for glucose levels during the ITT was significantly reduced only in WT EXE mice (**Fig. 4G**), as were plasma insulin levels (**Fig. S3A**) and the HOMA-IR and ADIPO-IR indices (**Fig. 4H,I**). Interestingly, in ADTRP KO mice, lower plasma insulin levels (**Fig. S3A**) and lower HOMA-IR and ADIPO-IR indices (**Fig. 4H,I**) were already observed in SED animals compared to WT animals; a similar pattern was observed for plasma cholesterol levels (**Table 1**). Subsequent correlation analysis showed that improvements in insulin sensitivity resulting from chronic exercise are closely linked to reductions in body fat (**Fig. 4J,K**; **Fig. S3B)**. The above results suggest that chronic exercise in HFD-fed obese mice improves insulin sensitivity, primarily in WT animals, as the lack of ADTRP hydrolase improves glucose homeostasis even in SED animals.

**Table 1.**
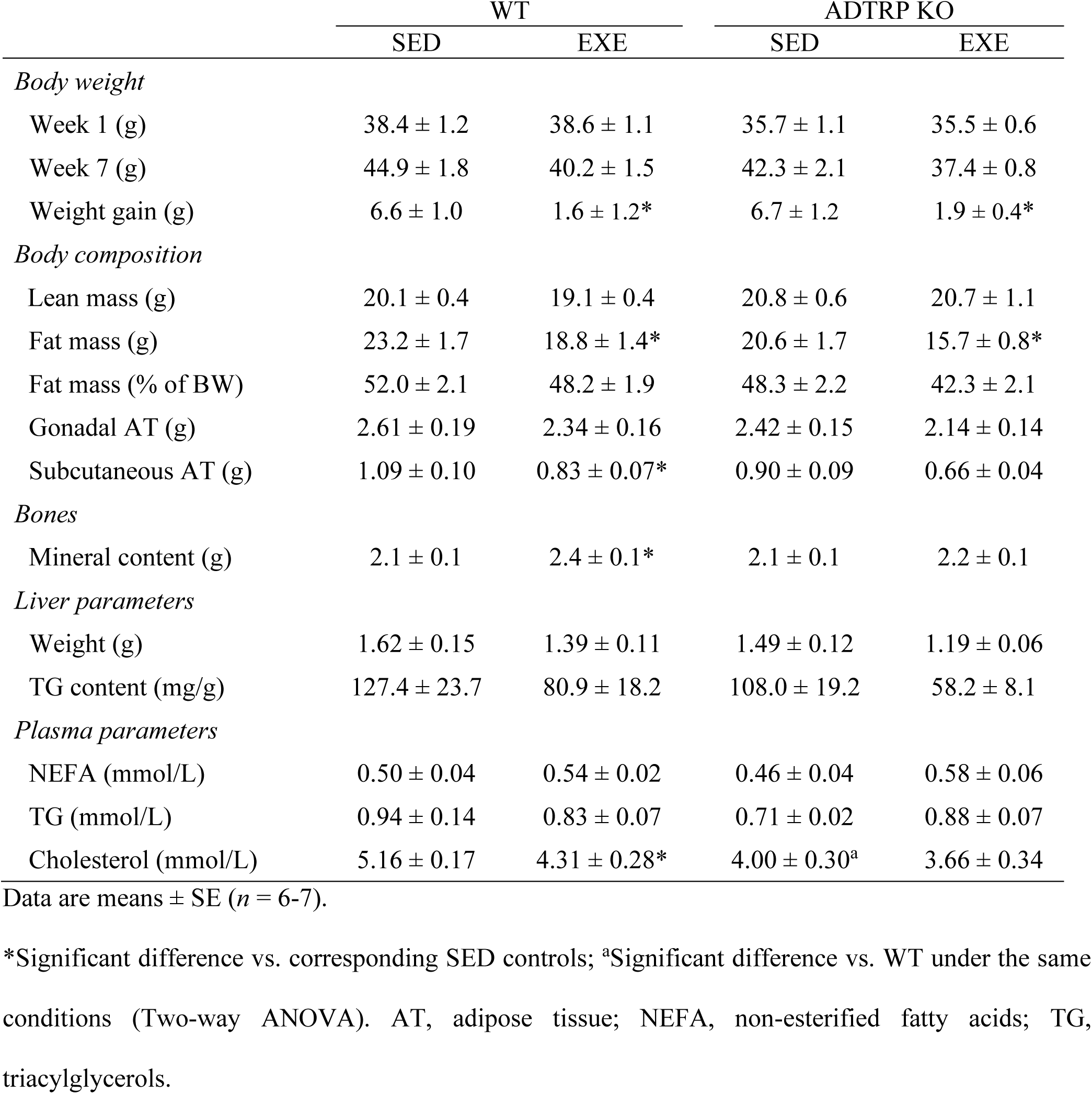
Body weight, body composition and metabolic parameters in WT and ADTRP KO mice subjected or not to chronic exercise intervention

### Chronic exercise elicited changes in gene expression consistent with improved AT function

To assess how acute exercise modulates AT quality, we first examined the mRNA levels of genes related to FAHFA metabolism and AT function, focusing on pathways such as *de novo* lipogenesis, lipolysis, and FAHFA degradation (**Fig. 5**). In gonadal AT, exercise increased the expression of *Slc2a4*, encoding the insulin-regulated glucose transporter GLUT4, in both WT and ADTRP KO mice, whereas *Dgat2*, *Srebf1*, and *Lipe*, i.e., genes associated with lipogenesis or lipolysis, were increased exclusively in WT mice, with no changes observed in ADTRP KO mice. On the other hand, exercise increased the expression levels of another lipogenic regulator, *Mlxipl*, exclusively in ADTRP KO mice (**Fig. 5A**). In addition, the expression levels of many genes related to lipogenesis and lipolysis showed a strong negative correlation with fat mass (**Fig. 5B**). Moreover, in WT EXE mice, the expression of the FAHFA-degrading enzyme androgen-induced gene 1 (*Aig1*) was significantly downregulated (**Fig. 5A**), and its expression generally showed a strong positive correlation with body fat mass (**Fig. 5B**). An effect of genotype was also observed, since ADTRP KO SED mice showed higher *Lipe* and lower *Aig1* levels than their WT counterparts (**Fig. 5A**). In subcutaneous AT, chronic exercise-induced changes in gene expression were less pronounced, although in ADTRP KO mice, increased expression of the *Slc2a4* was observed (**Fig. 5C**), and also a negative correlation with fat mass was noted (**Fig. 5D**). In summary, gene expression analysis indicated that regular exercise counteracts the detrimental effects of HFD feeding, leading to depot- and genotype-specific improvements in AT function.

**Figure 4.**
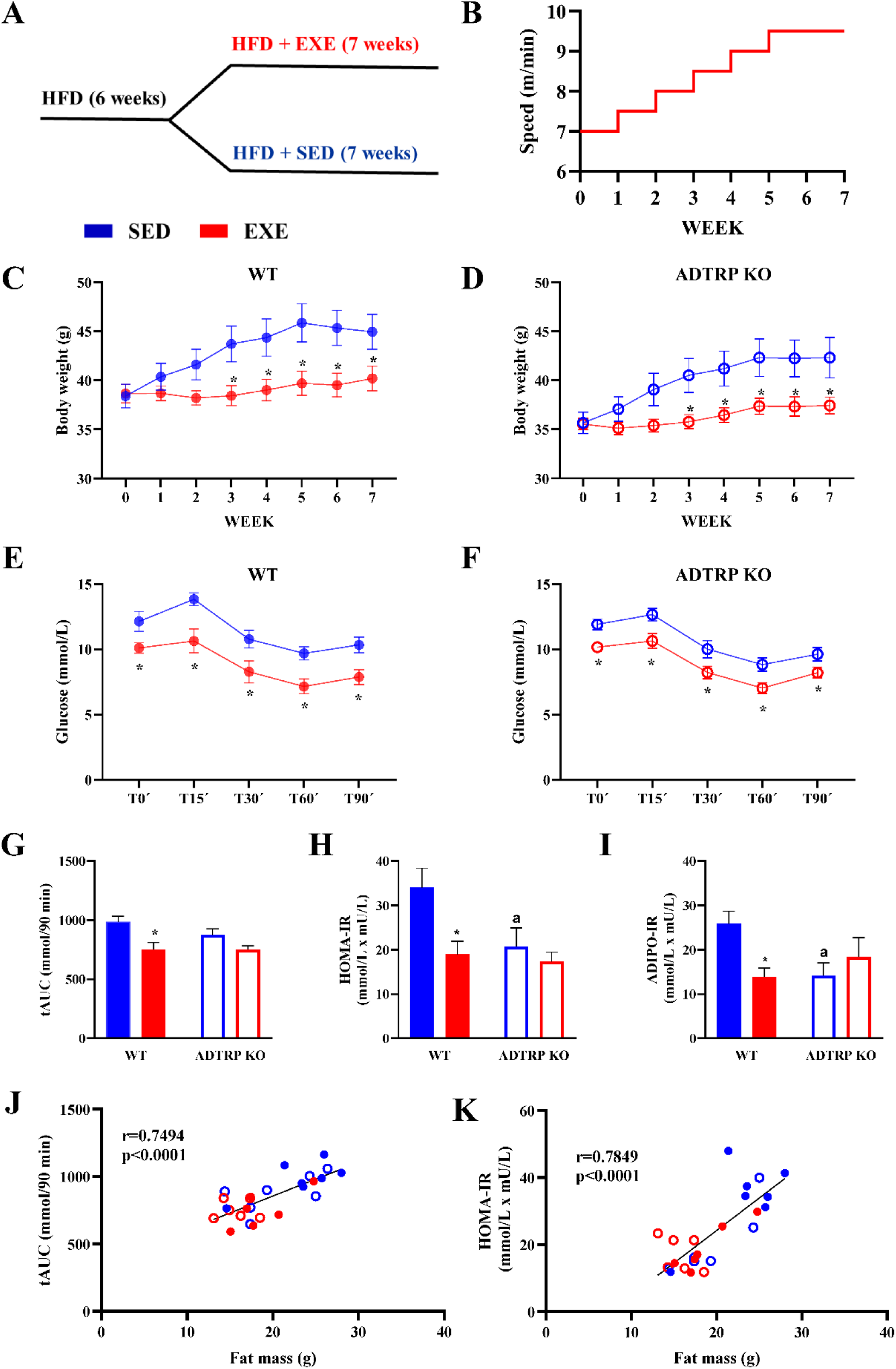
The effects of chronic exercise on body weight and glucose homeostasis in obese male WT and ADTRP KO mice fed HFD. A-B: Duration of HFD administration before and during the chronic exercise program (A) and treadmill exercise protocol during the long-term intervention (B). **C-D:** Effect of regular exercise on body weight in WT (C) and ADTRP KO (D) mice. **E-F:** Changes in blood glucose during the ITT in WT (E) and ADTRP KO (F) mice. **G-I:** Total AUC for glucose in the ITT (G), HOMA-IR index (H), and ADIPO-IR index (I) in WT and ADTRP KO mice. ^a,^*Significant difference vs. WT and SED, respectively; *p*≤0.05 (two-way ANOVA). **J-K:** Relationships between body fat mass and either the total AUC for glucose in the ITT (J), or the HOMA-IR index (K). The graphs show the Pearson correlation coefficients (r), and the statistical significance of the correlations.

**Figure 5.**
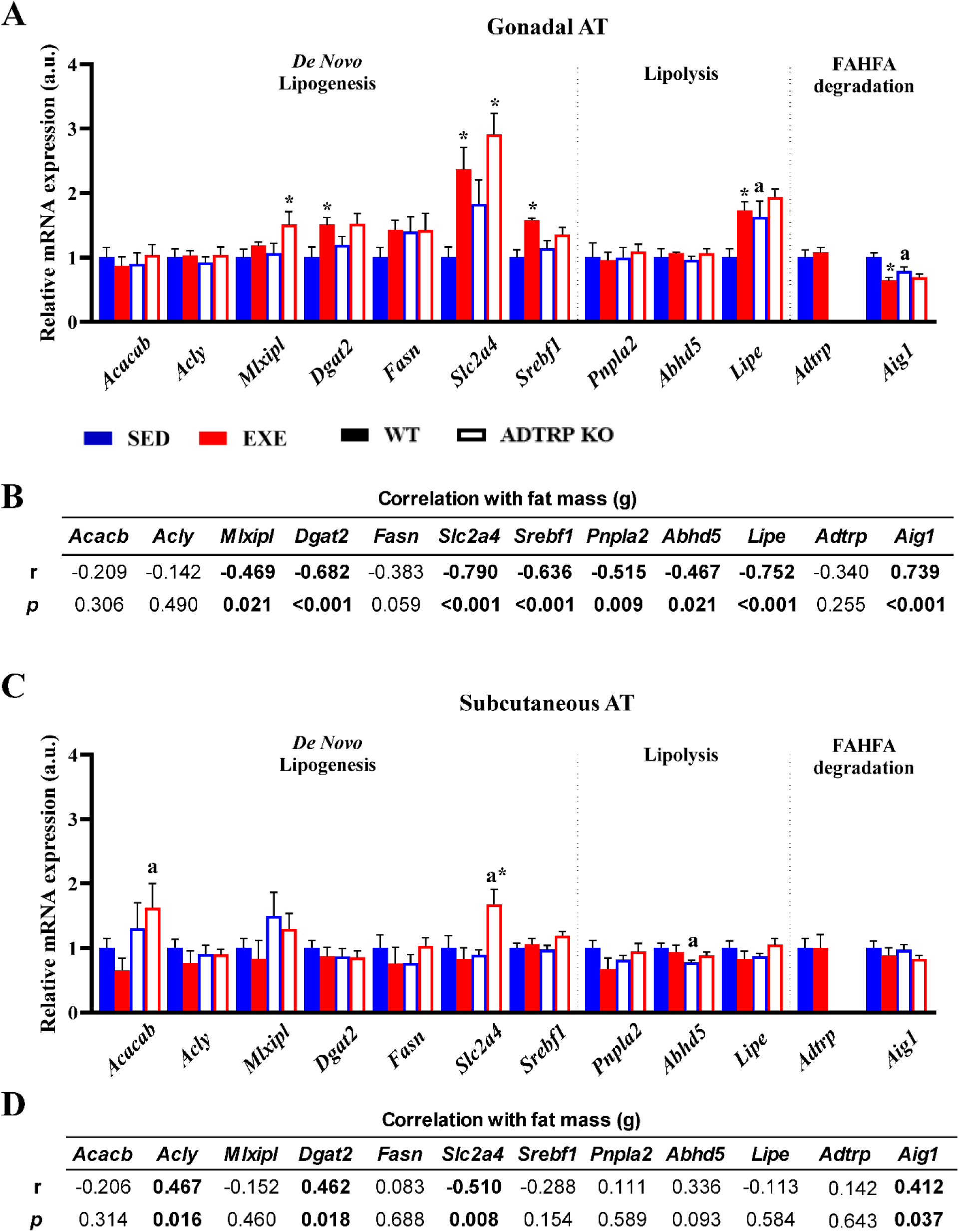
Changes in the expression of FAHFA metabolism-related genes in AT of obese mice fed HFD and subjected to chronic exercise. A: Expression of genes related to lipogenesis, lipolysis, and FAHFA degradation in gonadal AT of WT and ADTRP KO mice. **B:** Relationship between fat mass and expression levels of FAHFA metabolism-related genes in gonadal AT; Table shows Pearson correlation coefficients (r) and the statistical significance of the correlations. **C:** Expression of genes related to lipogenesis, lipolysis, and FAHFA degradation in subcutaneous AT of WT and ADTRP KO mice. **D:** Relationship between fat mass and expression levels of FAHFA metabolism-related genes in subcutaneous AT; Table shows the Pearson correlation coefficients (r) and the statistical significance of the correlations. ^a,^*Significant difference vs. WT and corresponding SED controls, respectively; *p*≤0.05 (two-way ANOVA).

### Chronic exercise induced a shift of FAHFA from the free form to TG EST in gonadal AT of WT mice fed ad libitum

In WT EXE mice, which were analyzed at the end of the intervention under *ad libitum* feeding conditions, total levels of free FAHFA and vast majority of detected FAHFA regioisomers in gonadal AT were reduced compared to their SED counterparts, whereas no such effect was observed in ADTRP KO EXE mice (**Fig. 6A,D**). In subcutaneous AT, ADTRP KO SED mice exhibited higher total levels of free FAHFA compared with WT SED mice, and exercise had no effect on total FAHFA levels or on the levels of individual FAHFA regioisomers, regardless of genotype (**Fig. 6B,E**). Plasma FAHFA levels were not affected by either genotype or chronic exercise (**Fig. 6C,F**). In SED mice, most of the detected FAHFA regioisomers followed the pattern as total FAHFA levels; i.e., in ADTRP KO mice, there were minimal changes in their levels in gonadal AT (**Fig. S4A**) and plasma (**Fig. S4C**), while the levels of a number of regioisomers were elevated in subcutaneous AT (**Fig. S4B**) compared with their WT counterparts. Overall, in HFD-fed obese mice, chronic exercise affected free FAHFA levels in gonadal AT, but not in subcutaneous AT or in plasma, whereas ADTRP deficiency primarily increased FAHFA levels in subcutaneous AT.

**Figure 6.**
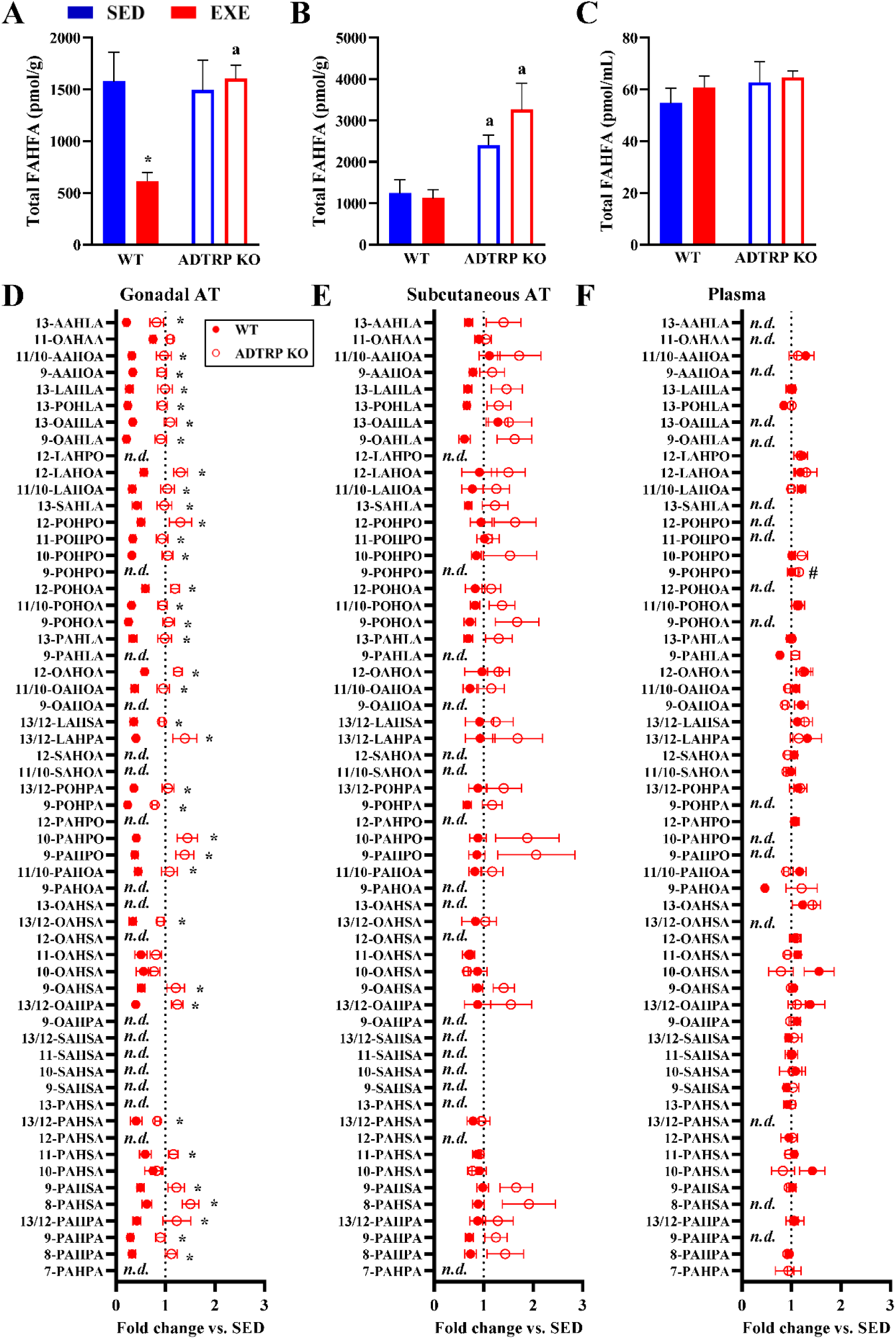
Free FAHFA levels in obese mice fed HFD and subjected to chronic exercise. A-C: Total free FAHFA levels in gonadal AT (A), subcutaneous AT (B), and plasma (C) of WT and ADTRP KO mice subjected or not to chronic exercise. ^a,^*Significant difference vs. WT and corresponding SED controls, respectively; *p*≤0.05 (two-way ANOVA). **D-F:** Levels of individual FAHFA regioisomers in gonadal AT (D), subcutaneous AT (E), and plasma (F) of WT and ADTRP KO mice subjected to chronic exercise; (fold change vs. corresponding SED controls). FAHFA species were sorted by total number of double bonds. *^,#^Significant difference vs. SED controls from the WT and ADTRP KO group, respectively; *p*≤0.05 (*t*-test).

To determine whether this local reduction in free FAHFA reflects a shift toward their esterified storage form (i.e., TG EST), we subsequently performed a lipidomic LC-MS analysis of gonadal AT. We focused specifically on FAHFA-containing TG EST and found that among 11 identified TG EST species, 7 were elevated in WT EXE mice (**Fig. 7A**). Several of these species contained FAHFA with palmitoleic acid esterified to a hydroxy-fatty acid (e.g., POHPA, POHOA; **Fig. 7A**). Regarding the effect of ADTRP deficiency alone, several TG species showed a trend toward increased levels compared with WT mice; however, these changes did not reach statistical significance (**Fig. S5**). Beyond TG EST, we identified 26 other lipid classes within the gonadal AT (**Fig. 7B**). The lipidomic profile of WT EXE mice indicated a functional shift toward lipid storage; specifically, we observed lower levels of acylcarnitines and lipolytic intermediates (free fatty acids and monoacylglycerols), along with an accumulation of triacylglycerol species (**Fig. 7B**). In contrast, in ADTRP KO EXE mice, the lipidomic profile of gonadal AT remained virtually unaffected relative to their SED counterparts (**Fig. 7B**). Given the distinct regulation of TG EST levels (elevated) and levels of either free FAHFA or lipolytic intermediates (decreased) in gonadal AT, specifically in WT EXE mice (**Fig. 7C**), our goal was also to explore the relationship between the above variables and the degree of insulin resistance. Therefore, correlation analyses between the AT levels of the aforementioned lipids and the HOMA-IR index were performed (**Fig. 7D**). In WT mice, TG EST levels did indeed exhibit a significant negative correlation with HOMA-IR, whereas levels of free FAHFA, free fatty acids, and monoacylglycerols were all positively correlated with HOMA-IR. In contrast, in ADTRP KO mice, only diacylglycerol levels showed a positive correlation with HOMA-IR (**Fig. 7D**). Overall, the above changes observed following chronic exercise demonstrate a metabolic shift in WT mice, in which TG EST levels correlate significantly with improved insulin sensitivity.

**Figure 7.**
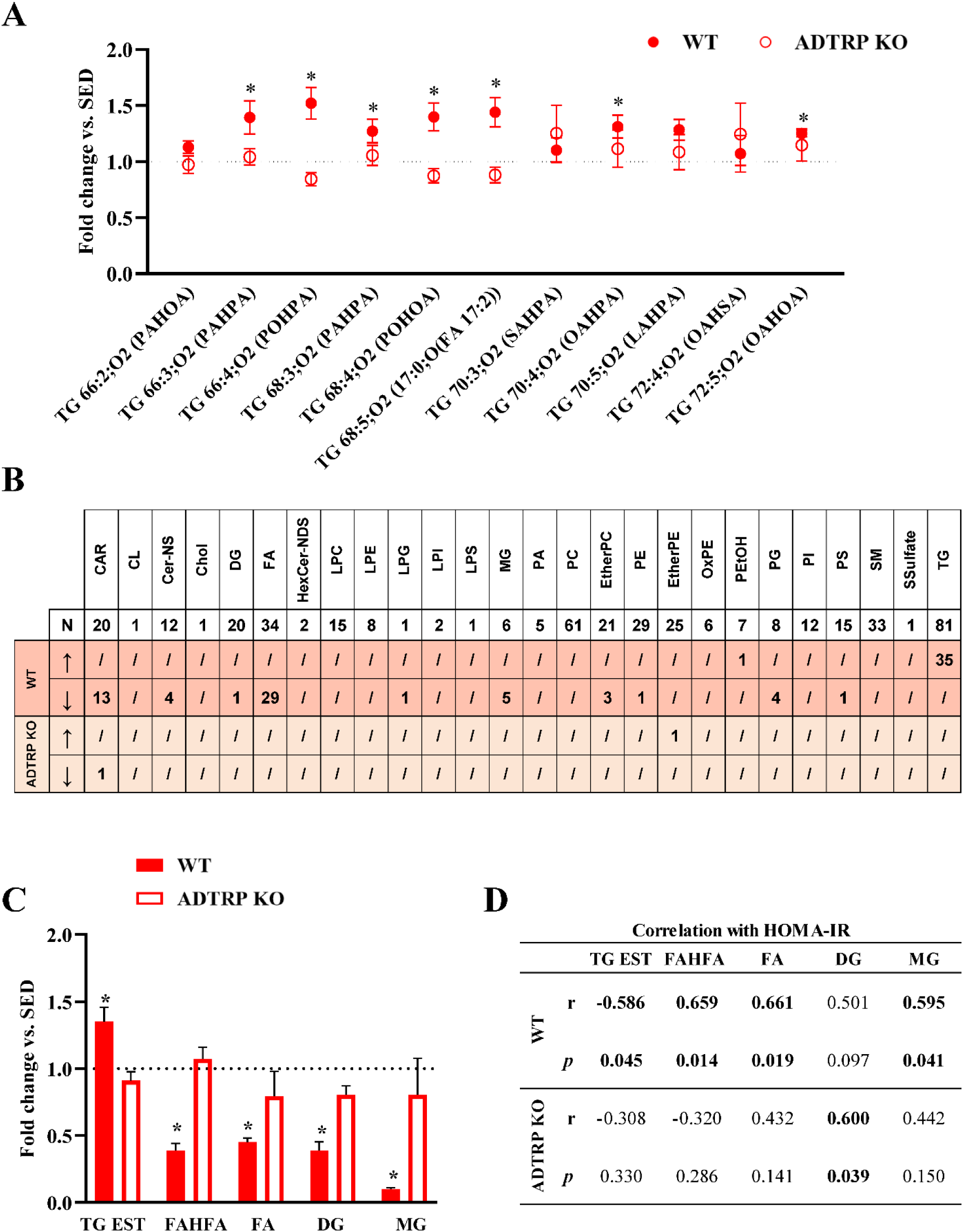
Lipidomic analysis of gonadal AT of obese mice fed HFD and subjected to chronic exercise. A: Levels of TG EST in WT and ADTRP KO mice subjected to exercise (fold change vs. corresponding SED controls). *^,#^Significant difference vs. SED controls from the WT and ADTRP KO group, respectively; *p*≤0.05 (*t*-test). **B:** An overview of the main lipid classes detected and the number of significantly altered lipid species within each class in EXE mice of both genotypes (vs. corresponding SED controls); *p*≤0.05 (*t*-test). **C:** Total levels of lipid species within the selected lipid classes (TG EST, free FAHFA, and lipid classes associated with lipolysis) in WT and ADTRP KO mice subjected to exercise (fold change vs. corresponding SED controls). *Significant difference vs. corresponding SED controls; *p*≤0.05 (*t*-test). **D:** Relationship between the degree of insulin resistance (i.e., HOMA-IR index) and the total levels of lipid species within the selected lipid classes; Table shows Pearson correlation coefficients (r) and the statistical significance of the correlations. CAR, Acylcarnitines; CL, Cardiolipins; Cer-NS, Ceramide non-hydroxyfatty acid-sphingosines; Chol, Cholesterol; DG, Diacylglycerols; FA, Free fatty acids; HexCer-NDS, Hexosylceramide non-hydroxyfatty acid-dihydrosphingosines; LPC, Lysophosphatidylcholines; LPE, Lysophosphatidylethanolamines; LPG, Lysophosphatidylglycerols; LPI, Lysophosphatidylinositols; LPS, Lysophosphatidylserines; MG, Monoacylglycerols; PA, Phosphatidic acid; PC, Phosphatidylcholines; EtherPC, Ether-linked phosphatidylcholines; PE, Phosphatidylethanolamines; EtherPE, Ether-linked phosphatidylethanolamines; OxPE, Oxidized phosphatidylethanolamines; PEtOH, Phosphatidylethanols; PG, Phosphatidylglycerols; PI, Phosphatidylinositols; PS, Phosphatidylserines; SM, Sphingomyelin; SSulfate, Sterol sulfates; TG, Triacylglycerols.

## Discussion

This study was designed in two complementary parts to delineate FAHFA metabolism in the context of acute and chronic exercise, while accounting for the influence of sex, genotype and diet-induced obesity. In the first part, we investigated sex differences in the response to acute exercise, focusing on lipolysis and FAHFA release in AT of WT mice and those with constitutively elevated FAHFA levels in AT (i.e., ADTRP KO mice).

Although no differences were observed in running time to exhaustion or in post-exercise lactate levels, suggesting a similar relative exercise intensity across all groups (29), acute exercise-induced lipolysis in AT was more pronounced in male compared to female mice. This finding is consistent with previous studies reporting sex differences in lipolytic responses (30–33). With regard to free FAHFA levels in AT, it has been shown that they increase following *in vitro* lipolysis activation (13), as well as in response to *in vivo* stimuli such as fasting (7, 14, 16, 17). Once liberated, FAHFA species can bind to their receptors and modulate the inflammatory response and insulin action (34). In our study, a single bout of exercise led to enhanced FAHFA release from AT in males, consistent with the more pronounced lipolytic response to acute exercise observed in male compared to female mice. Interestingly, this effect was observed only in a relatively small subset of FAHFA regioisomers. The mechanisms governing the transport of FAHFA across the cell membrane remain largely unexplored. It is therefore possible that certain FAHFA regioisomers are liberated during exercise-induced lipolysis but are not efficiently transported out of AT. This interpretation is supported by previous studies, demonstrating that fasting leads to an increase in free PAHSA levels in AT without a corresponding rise in circulating FAHFA levels (7).

Adipose triglyceride lipase has been described as a biosynthetic enzyme for FAHFA (14, 35), with fasting reported to stimulate both its lipolytic and transacylase activities involved in *de novo* FAHFA synthesis (17). In contrast, HSL, another principal lipase, has been identified as an FAHFA hydrolase involved in FAHFA degradation in AT (14). Thus, upon activation, HSL may simultaneously liberate FAHFA from TG EST and promote their degradation, thereby potentially limiting their availability for receptor binding and downstream physiological effects. Interestingly, in our study, *ex vivo* inhibition of HSL activity in AT explants isolated from mice following acute exercise revealed sex-dependent differences in HSL’s involvement in FAHFA metabolism.

Specifically, the ability of activated HSL to degrade FAHFA appeared to be more pronounced in females than in males, despite lower lipolytic activity in the former. Notably, this pattern was observed across most of the identified FAHFA regioisomers and was consistent in AT explants from both WT and ADTRP KO mice, strengthening the relevance of this finding. Our results point to a potentially important sex-specific mechanism regulating FAHFA metabolism, based on differential involvement of the enzymes ATGL and HSL during activated lipolysis. Therefore, further research into the role of HSL in FAHFA dynamics under various metabolic conditions is warranted.

Based on the greater acute exercise-induced lipolysis and higher FAHFA release from AT observed in male mice, we designed a chronic exercise intervention in obese HFD-fed males. The aim was to determine whether regular exercise could improve AT function and FAHFA metabolism while alleviating the adverse metabolic effects of HFD administration, and whether the effect of exercise could be potentiated under conditions of persistently elevated FAHFA levels in AT. Some human studies indicate a possible link between metabolic benefits and greater exercise intensity (36); however, during low-intensity exercise, lipid utilization predominates, with NEFA derived from AT serving as the primary fuel for skeletal muscle (37, 38), while AT lipolysis does not appear to increase further as exercise intensity rises (39). Our study, therefore, focused on lower-intensity exercise training, both because of its potential to induce greater lipolytic activity and because this type of exercise intervention is easier to implement in obese mice.

Chronic exercise in our experimental model suppressed body weight gain on HFD and improved insulin sensitivity. The beneficial effects of regular exercise on body weight regulation and insulin sensitivity are well documented in the literature (40–43). Indeed, correlation analyses indicated that these exercise-induced improvements in glucose homeostasis and insulin sensitivity were closely associated with reductions in fat mass. Regarding the effect of chronic exercise, the observed reduction in free FAHFA levels in gonadal AT of WT mice was not expected, particularly given the increased expression of genes associated with FAHFA synthesis and improved AT health, along with reduced expression of *Aig1*, i.e., a FAHFA hydrolase associated with chronic inflammation (44), in response to chronic exercise. However, it is important to note that the timing of tissue sampling may have influenced the measured FAHFA levels, as free FAHFA levels are tightly regulated by fasting-feeding cycles (17). In our study, insulin sensitivity improved in WT mice in response to chronic exercise, so that lipolysis should be effectively suppressed at the level of AT in animals dissected in a fed state. Indeed, using LC-MS-based lipidomic analysis, we found lower levels of lipolytic products, including free fatty acids, in AT of trained WT mice compared with insulin-resistant sedentary controls. To further elucidate the effect of chronic exercise on FAHFA metabolism, we therefore analyzed the content and composition of TG EST, a class of TG species containing FAHFA esterified to glycerol, in AT. In contrast to free FAHFA, TG EST levels increased in WT mice in response to exercise. This finding is consistent with our previous observation in older women, in whom exercise training was linked to improved AT function, enhanced insulin sensitivity, and increased FAHFA synthesis (22). Notably, several TG EST species elevated by chronic exercise contained palmitoleic acid (16:1*n*-7), a fatty acid independently associated with improved insulin sensitivity (45). Furthermore, correlation analyses indicated a negative relationship between TG EST levels in AT and the degree of insulin resistance, as measured by the HOMA-IR and AdipoIR indices; this further supports a mechanism in which improved insulin sensitivity in AT of exercising WT mice enabled more effective suppression of lipolysis in the fed state. On the other hand, the lack of increase in TG EST in AT of ADTRP-deficient mice following long-term exercise could be explained by the fact that the absence of functional ADTRP confers certain metabolic advantages on these animals independent of exercise, as evidenced by lower body weight and reduced body fat mass compared with their WT counterparts, although these differences did not reach statistical significance due to considerable intra-group variability. It is therefore possible that the improved baseline metabolic phenotype of ADTRP-deficient mice limited the positive effects of regular exercise on TG EST levels and overall AT health.

In summary, this study provides novel insights into the regulation of FAHFA metabolism by exercise and sex. Acute exercise affected the release of free FAHFA from AT in a sex-dependent manner, specifically in the context of stimulated lipolysis and HSL-dependent FAHFA degradation, which was predominant in females. In addition, chronic exercise in male mice improved insulin sensitivity of AT and increased tissue levels of TG EST, which negatively correlated with the degree of whole-body insulin resistance, suggesting a possible link between FAHFA metabolism in AT and improvements in whole-body metabolism. The precise role of HSL in FAHFA metabolism during activated lipolysis, as well as the sex-specific effects of long-term exercise on FAHFA regulation and AT health, will need to be clarified in future studies.

## Declarations

### Ethics approval and consent to participate

Animal experiments were approved by the Institutional Animal Care and Use Committee and the Committee for Animal Protection of the Czech Academy of Sciences (Approval Number: 62/2023), following the EU Directive 2010/63/EU on protecting animals used for scientific purposes.

## Consent for publication

Not applicable.

## Availability of data and materials

The datasets analyzed during the current study are available from the corresponding author on reasonable request.

## Authors’ contributions

MM contributed to data collection and analysis, interpretation of results, and drafted the manuscript; OH participated in the study design, contributed to data collection and analysis, and interpretation of results; MRi contributed to data analysis; VK and PZ contributed to data collection and analysis; TC contributed to data collection and analysis; LR participated in the study design and interpretation of results; OK contributed to data analysis; MRo obtained funding, participated in the study design, contributed to data collection and interpretation of results, and drafted the manuscript. All authors contributed to the editing of the manuscript and approved the final version.

## Acknowledgments

We would like to thank Alan Saghatelian (Salk Institute for Biological Studies, La Jolla, CA, USA) for providing breeding pairs of ADTRP KO mice.

## List of abbreviations

ADTRP, androgen-dependent tissue factor pathway inhibitor regulating protein; ADTRP KO, ADTRP knockout; AIG1, androgen-induced gene 1; AT, adipose tissue; FAHFA, fatty acid esters of hydroxy fatty acids; HFD, high-fat diet; HOMA-IR, homeostatic model assessment of insulin resistance; HSL, hormone-sensitive lipase; NEFA, non-esterified fatty acids; PAHSA, palmitic acid esters of hydroxy stearic acids; TG EST, triacylglycerol estolide.

