## Supplementary information for "Higher baseline levels of fatty acid esters of hydroxy fatty acids do not further enhance the stimulatory effect of regular exercise on insulin sensitivity in obese mice"

**Methods**

*LC-MS analysis*

Lipidomic profiling of gonadal AT samples was performed using an untargeted workflow for the lipidome, metabolome, and exposome analysis, as reported before [1, 2]. Briefly, AT samples were extracted using a biphasic solvent system of cold methanol, methyl *tert*-butyl ether (MTBE), and 10% methanol. Then, LC-MS-based lipidomic profiling of (i) high-abundant TGs in positive ion mode and (ii) minor polar lipids in positive and negative ion modes was performed using the reversed-phase LC mechanism. Hydrophilic interaction chromatography (HILIC) in positive mode was used to analyze short-chain acylcarnitines. The LC-MS systems consisted of (i) a Vanquish UHPLC system (Thermo Fisher Scientific), an OptaMax NG ion source with a HESI-II probe (Thermo Fisher Scientific), and an Orbitrap Exploris 480 mass spectrometer (Thermo Fisher Scientific).

The YMC-Triart C18 ExRS column (150 mm × 2 mm i.d.; 1.9 μm particle size; YMC America, Devens, MA, USA) equipped with a holder-cartridge assembly with a C18 guard cartridge (Phenomenex) was utilized to separate FAHFA species. The separation was carried out at a flow rate of 0.2 mL/min, with the column maintained at 50°C. The mobile phase included (A) 70:30 water/acetonitrile with 0.1% acetic acid and (B) 50:50 isopropanol/acetonitrile with 0.1% acetic acid. Separation was achieved through the following gradient: 0–1 min 0% (B); 1–5 min from 0% to 90% (B); 5–23 min 90% (B), 23–24 min from 90% to 100% (B), 24–26 min 100% (B), 26–28 min from 100% to 0% (B), 28–31 min 0% (B) + 1 min preinjection steps. The injection volume was 10 μL. The sample temperature was kept at 4°C.

The ion source parameters were as follows: sheath gas pressure, 45 arbitrary units; aux gas flow, 10 arbitrary units; sweep gas flow, 2 arbitrary units; ion transfer tube temperature, 300°C; vaporizer temperature, 400°C; spray voltage: −2.5 kV. The mass spectrometer settings were set to MS1 mass range, *m*/*z* 505–610; MS1 resolving power, 90,000 FWHM (*m*/*z* 200); the number of data-dependent scans per cycle, 5; MS/MS resolving power, 15,000 FWHM (*m*/*z* 200). For MS/MS experiments, normalized collision energies of 20 and 30% were set up.

Samples were randomized within the sequence, with regular injections of quality control (QC) pool samples at the beginning, end, and every 10 actual samples. The sequence also included method blanks and serial dilution samples prepared from QC sample (0, 1/16, 1/8, 1/4, 1/2, 1).

The LC-MS instrumental files were processed using MS-DIAL v. 4.9.221218 software [3], including annotation of FAHFA using MS1 and MS/MS spectra. FAHFA were reported at the species level as the total number of carbon atoms and double bonds (e.g., FAHFA 34:0;O) and at the molecular species level (FAHFA 16:0/18:0;O). Exported data sets as signal intensities from the detector (peak heights) were further filtered by method blanks, a dilution series of QC sample, and QC samples (a relative standard deviation (RSD) >30%). Data were then normalized using locally estimated scatterplot smoothing (LOESS), the amount taken for the analysis, and using [^13^C_4_]-9-PAHSA internal standard. The subsequent regioisomer annotation of FAHFA was based on the previous run of authentic standards [4].

*FAHFA analysis*

Samples of AT (40–50 mg) and plasma (150 μL) were extracted according to previously described methods, with several modifications to improve extraction yield and FAHFA detection during LC-MS analysis [4, 5]. Samples were homogenized using a ball mill (see Section 2.8 of the main article) in a mixture of citric acid buffer and ethyl acetate (ratio 1:2, v/v) containing 1 ng of [^13^C_4_]-9-PAHSA (Cayman Chemical; Ann Arbor, MI, USA) as an internal standard. The ethyl acetate phase was collected and dried using a Savant SpeedVac (Thermo Fisher Scientific, Bremen, Germany). Based on optimization of extraction yield, we dissolved the dried extracts in 300 μL (200 μL for plasma) of chloroform instead of a mixture of ethyl acetate and hexane (5:95, v/v) and loaded the samples onto HyperSep SPE columns (500 mg/10 mL, 40–60 μm, 70 Å; Thermo Scientific) for further purification by solid-phase extraction (SPE). FAHFA were eluted with 5 mL ethyl acetate and dried. Before analysis, purified extracts were dissolved in 50 μL of a methanol/water mixture (95:5, v/v) to eliminate TG, potentially suppressing FAHFA signal. Samples were analyzed by LC-MS as before [4].

Table S1

Gene names and sequences of the oligonucleotide primers.

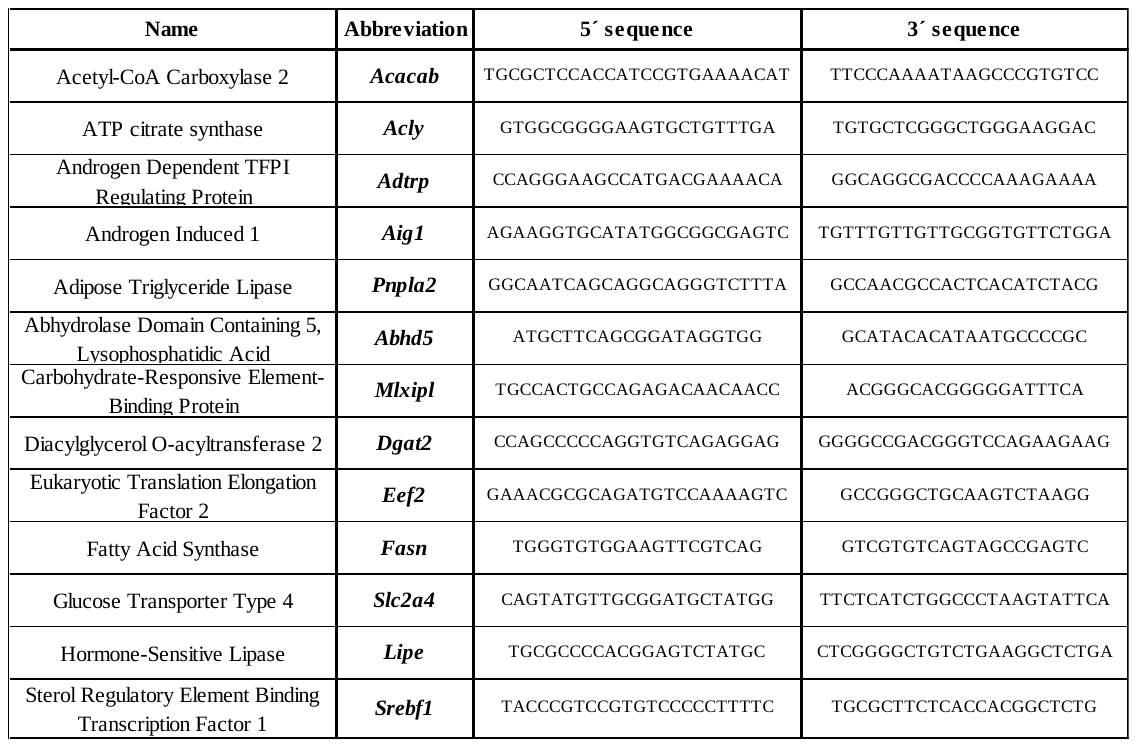

Table S2

List of annotated FAHFA in medium.

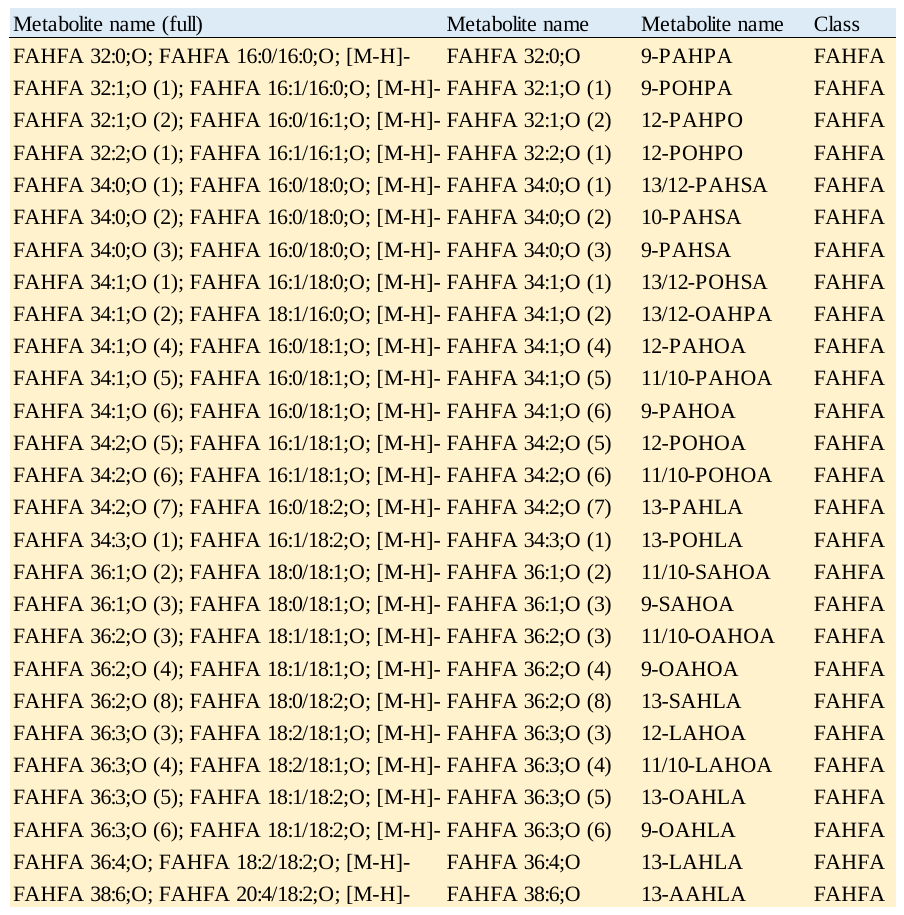

Table S3

List of annotated FAHFA in adipose tissues.

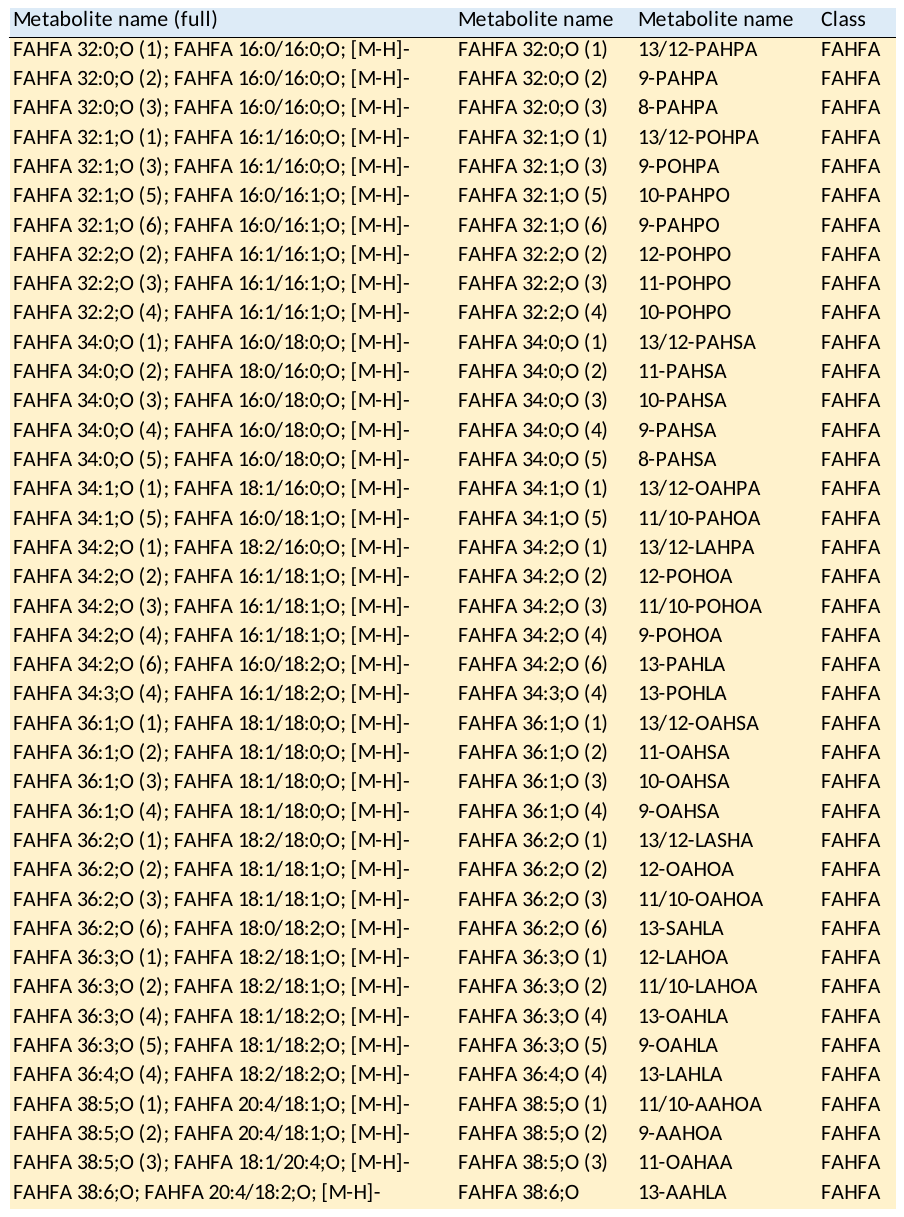

Table S4

List of annotated FAHFA in plasma.

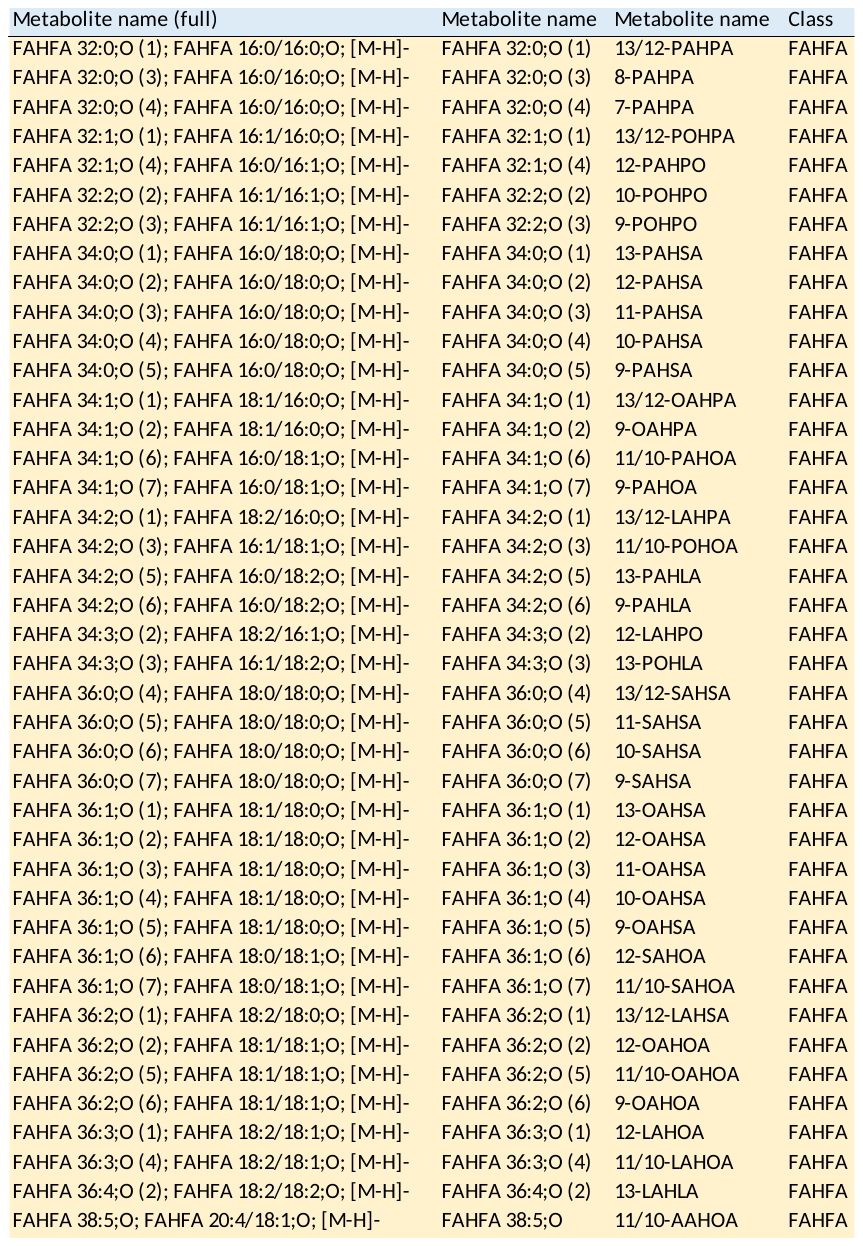

Table S5

List of annotated lipids.

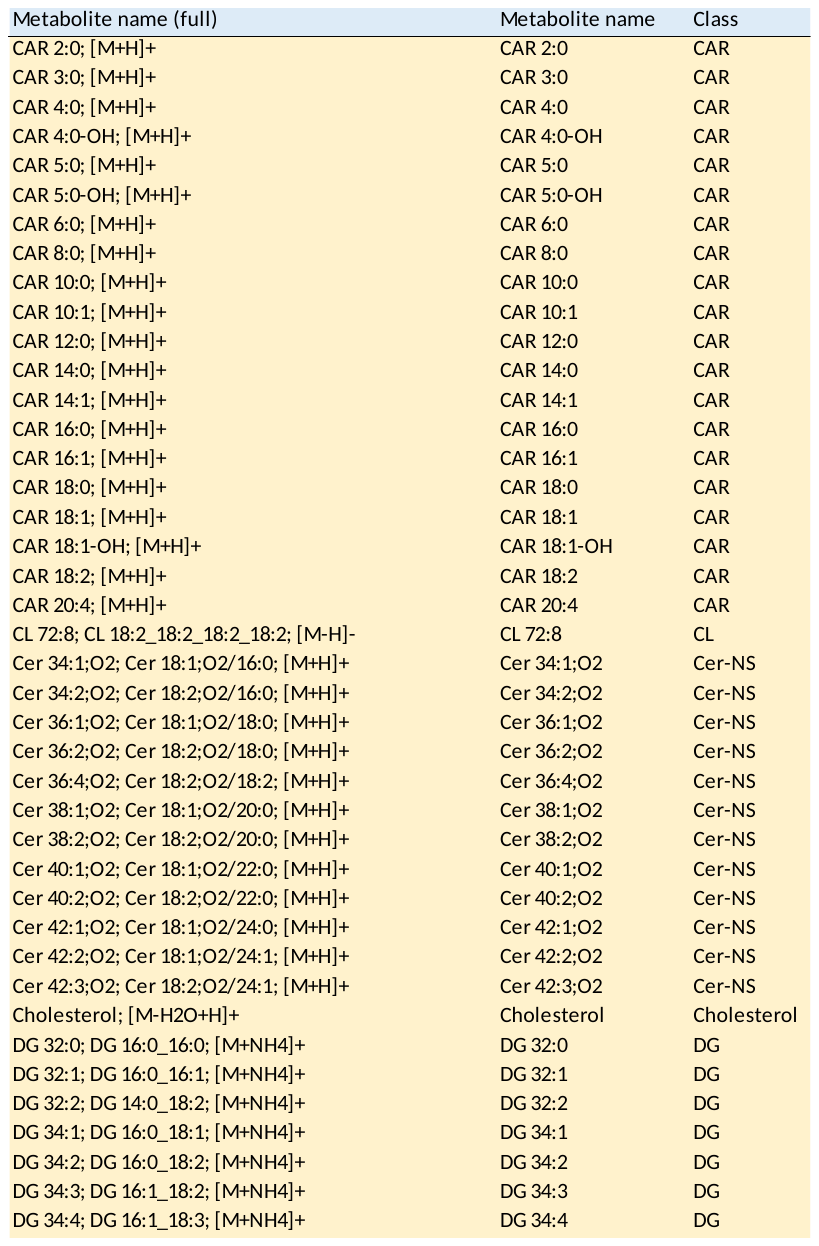

Table S5 (cont.)

List of annotated lipids.

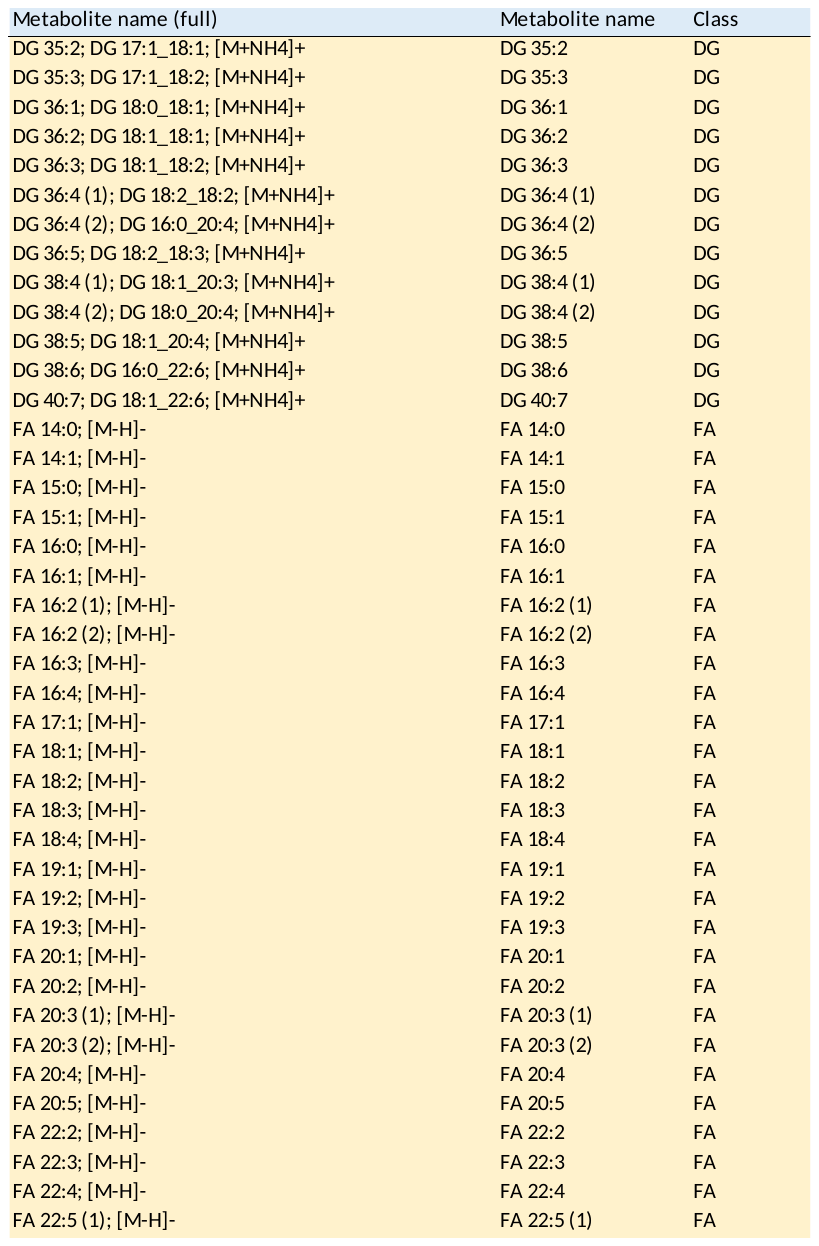

Table S5 (cont.)

List of annotated lipids.

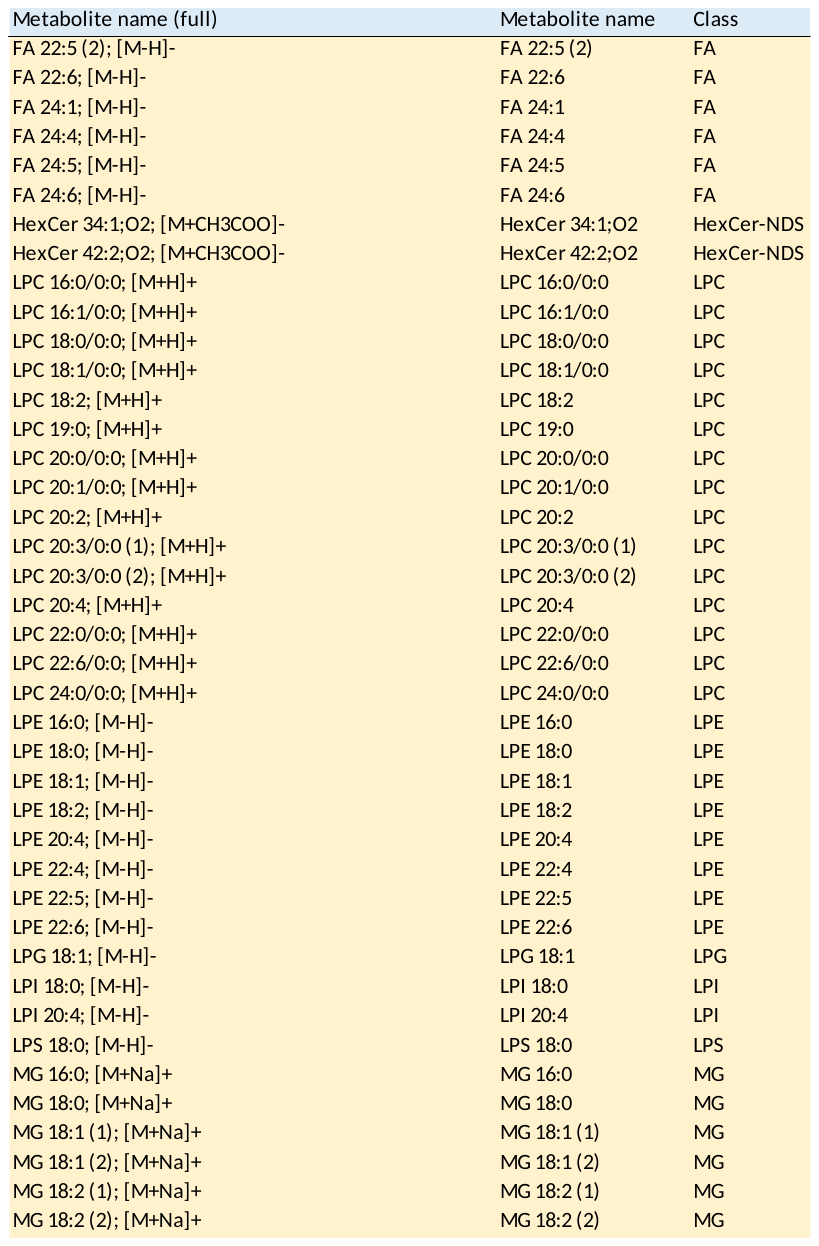

Table S5 (cont.)

List of annotated lipids.

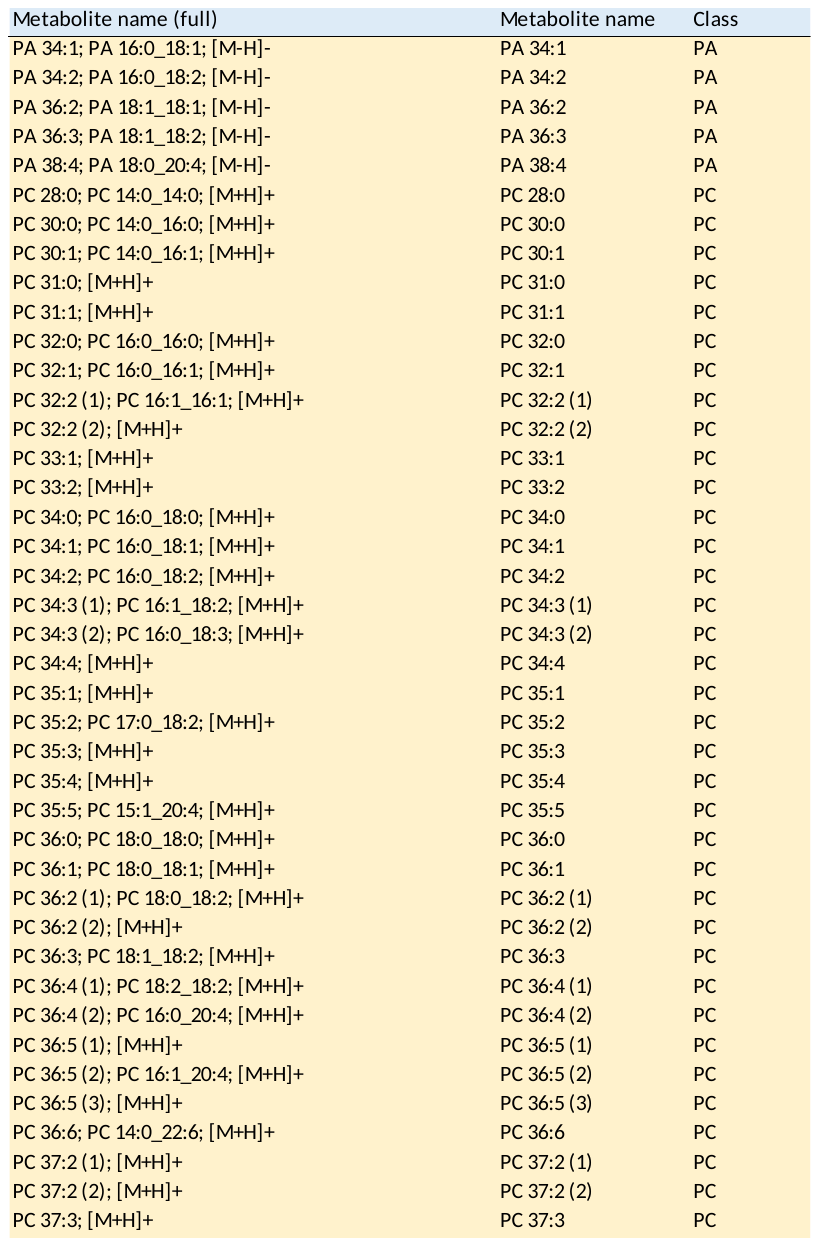

Table S5 (cont.)

List of annotated lipids.

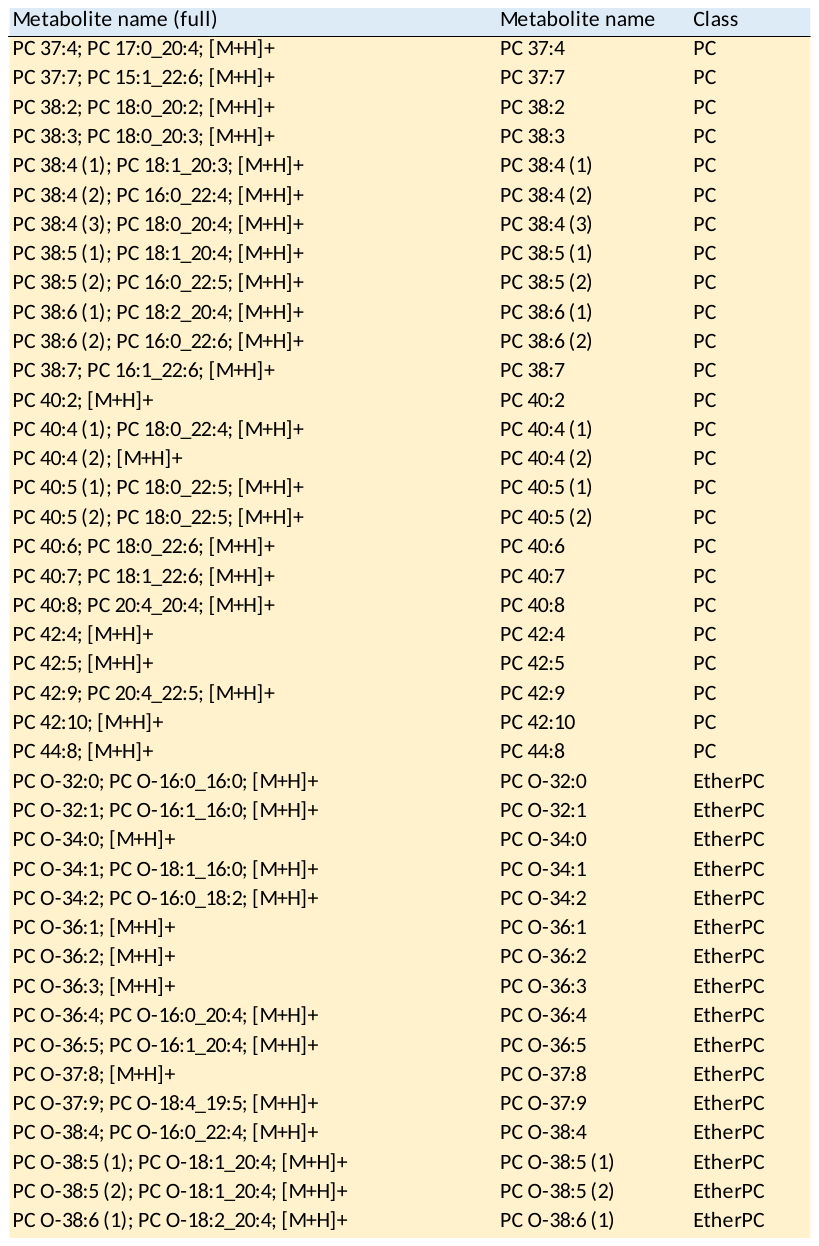

Table S5 (cont.)

List of annotated lipids.

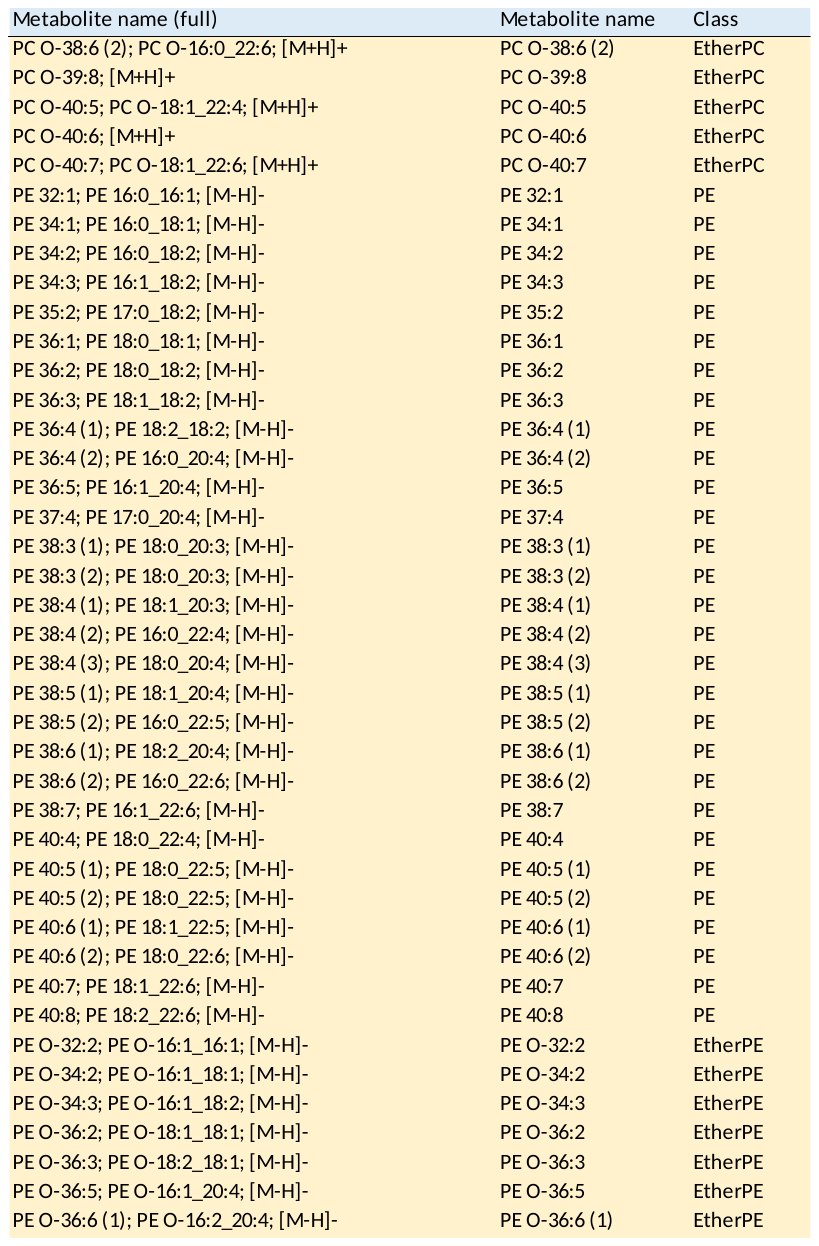

Table S5 (cont.)

List of annotated lipids.

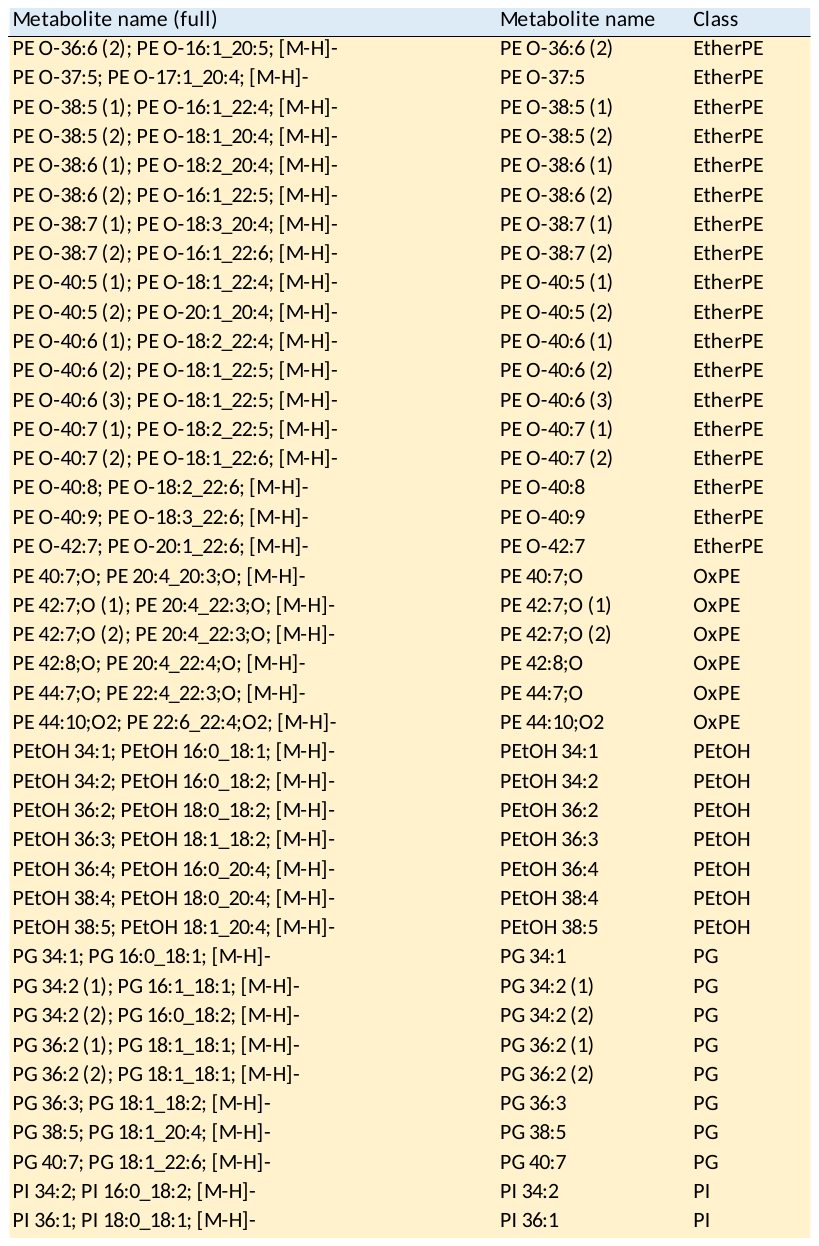

Table S5 (cont.)

List of annotated lipids.

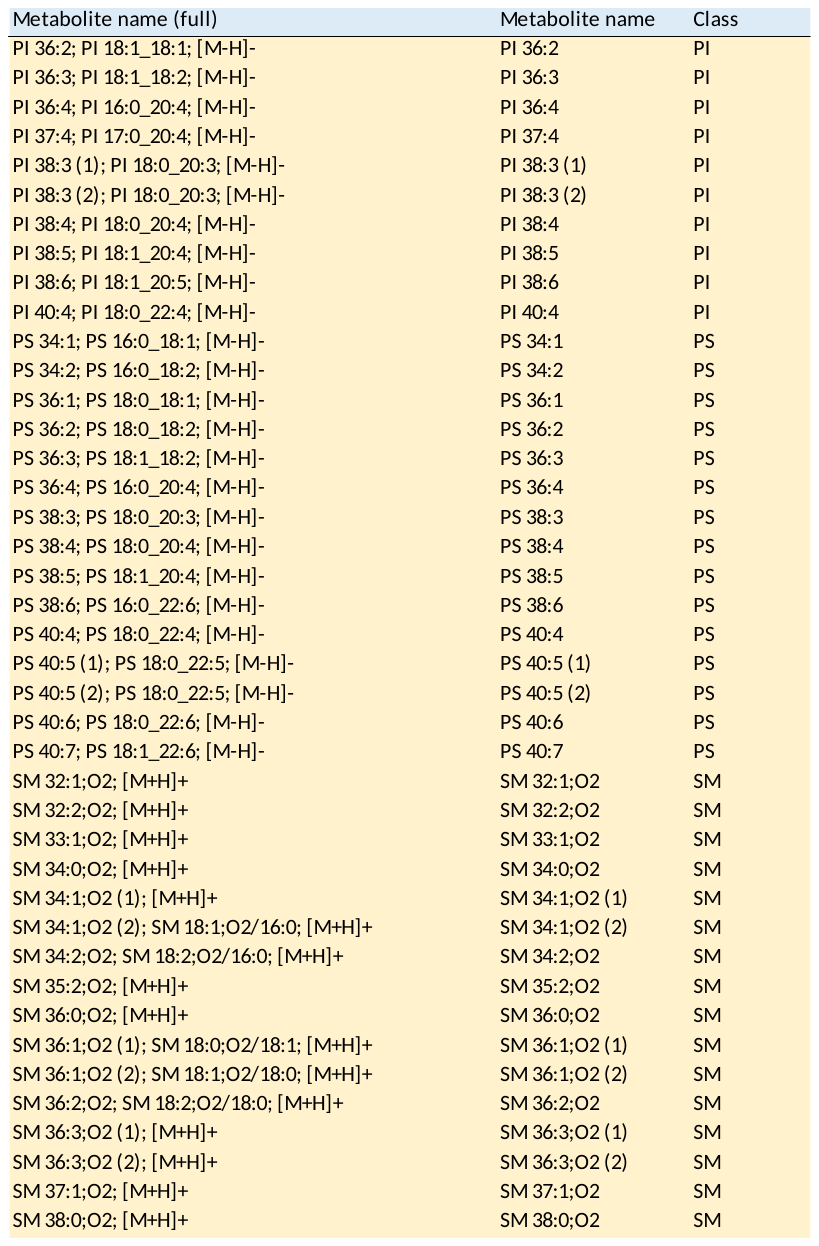

Table S5 (cont.)

List of annotated lipids.

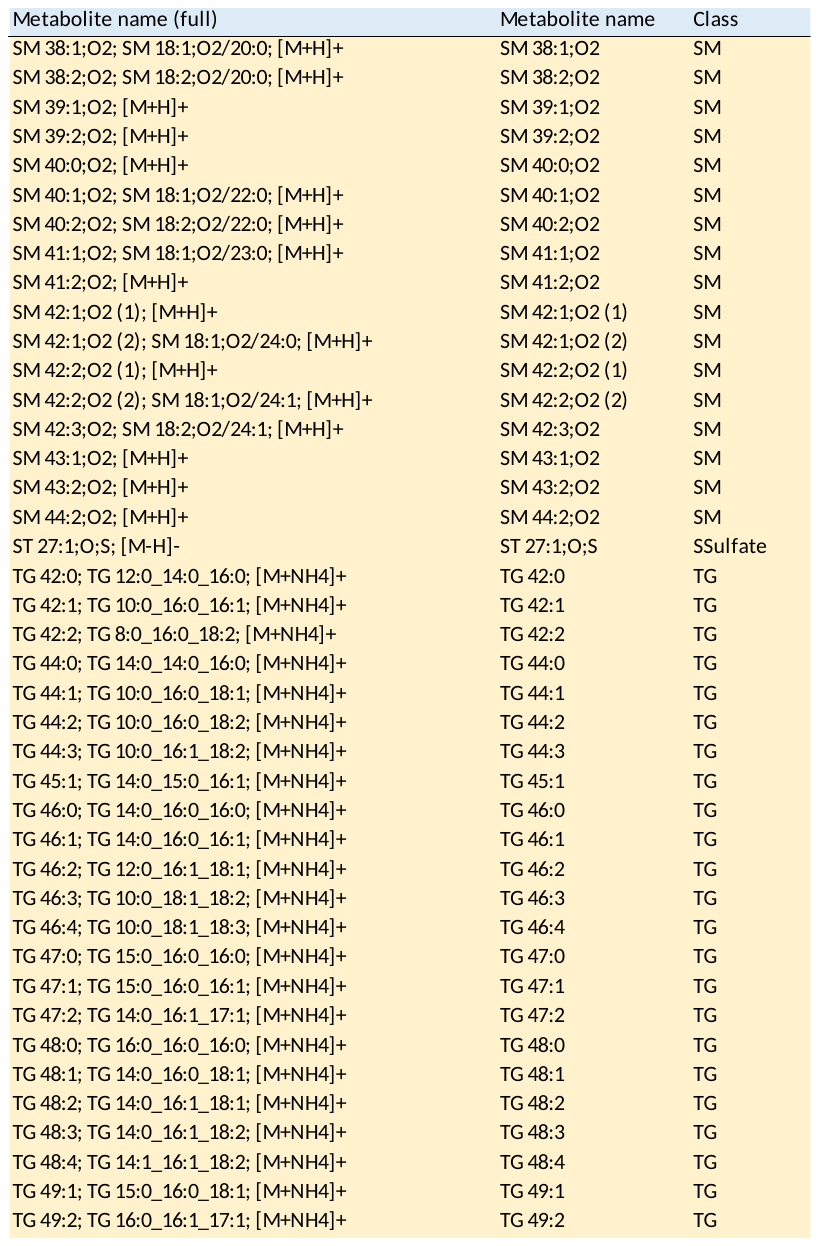

Table S5 (cont.)

List of annotated lipids.

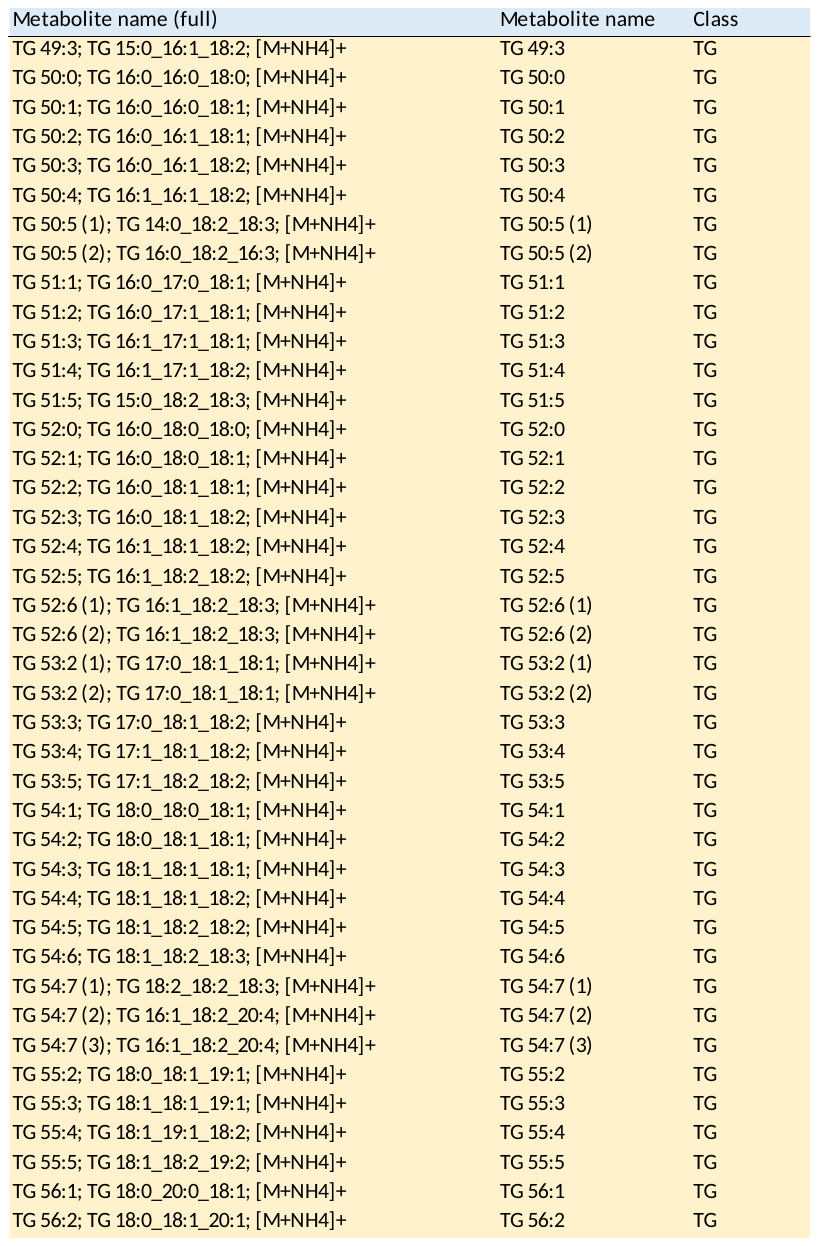

Table S5 (cont.)

List of annotated lipids.

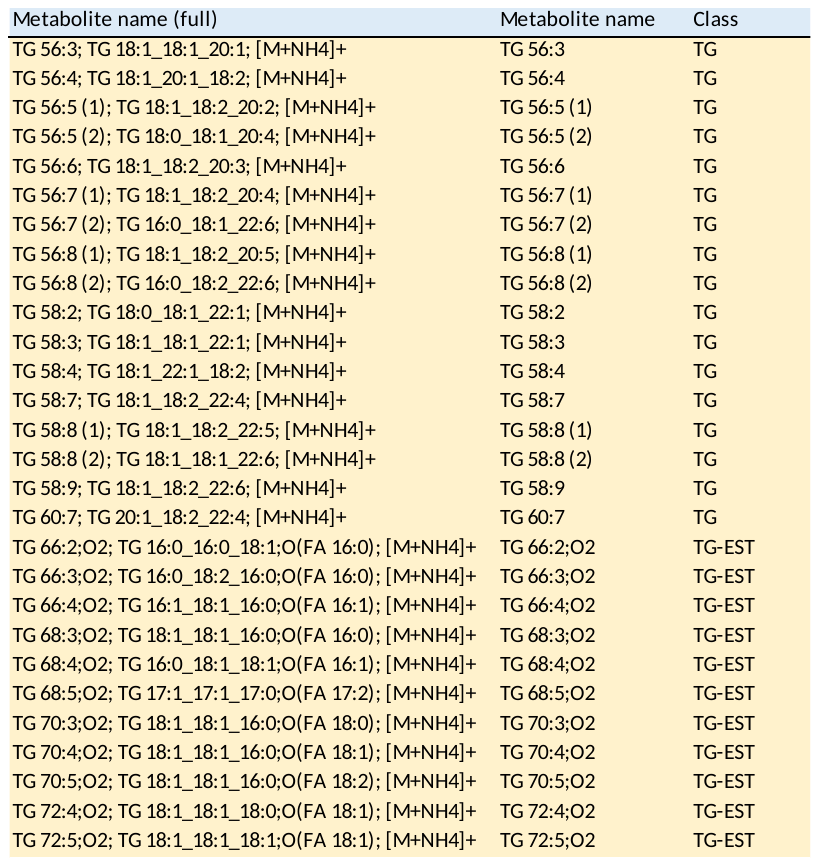

Table S6. Body weight, adiposity and metabolic parameters in male and female WT mice subjected or not to an acute bout of exercise

|  | WT | | | | | | |
| --- | --- | --- | --- | --- | --- | --- | --- |
|  | FEMALE | | |  | MALE | | |
|  | CON |  | EXE |  | CON |  | EXE |
| Body weight (g) | 22.9 ± 0.5 |  | 22.8 ± 0.5 |  | 32.3 ± 0.7^a^ |  | 32.0 ± 0.6^a^ |
| Gonadal AT (g) | 0.26 ± 0.04 |  | 0.21 ± 0.02 |  | 0.68 ± 0.11^a^ |  | 0.49 ± 0.05^a^ |
| Subcutaneous AT (g) | 0.12 ± 0.01 |  | 0.17 ± 0.01* |  | 0.19 ± 0.03^a^ |  | 0.22 ± 0.02 |
| Glucose (mmol/L) | 7.2 ± 0.3 |  | 5.3 ± 0.1* |  | 7.7 ± 0.2 |  | 5.7 ± 0.2* |

Data are means ± SE (*n*=6-7).

*, Statistically significant vs. CON and; ^a^, Statistically significant vs. FEMALE; *p* ≤ 0.05 (Two-way ANOVA).

|  | ADTRP KO | | | | | | |
| --- | --- | --- | --- | --- | --- | --- | --- |
|  | FEMALE | | |  | MALE | | |
|  | CON |  | EXE |  | CON |  | EXE |
| Body weight (g) | 22.8 ± 0.3 |  | 23.0 ± 0.5 |  | 31.9 ± 0.5^a^ |  | 31.6 ± 0.7^a^ |
| Gonadal AT (g) | 0.18 ± 0.02 |  | 0.19 ± 0.02 |  | 0.60 ± 0.11^a^ |  | 0.66 ± 0.13^a^ |
| Subcutaneous AT (g) | 0.10 ± 0.02 |  | 0.16 ± 0.01* |  | 0.17 ± 0.03 |  | 0.26 ± 0.05* |
| Glucose (mmol/L) | 6.9 ± 0.3 |  | 5.3 ± 0.3* |  | 7.8 ± 0.4 |  | 5.6 ± 0.4* |

Table S7. Body weight, adiposity and metabolic parameters in male and female ADTRP KO mice subjected or not to an acute bout of exercise

Data are means ± SE (n=6-7).

*, Statistically significant vs. CON and; ^a^, Statistically significant vs. FEMALE; *p* ≤ 0.05 (Two-way ANOVA).

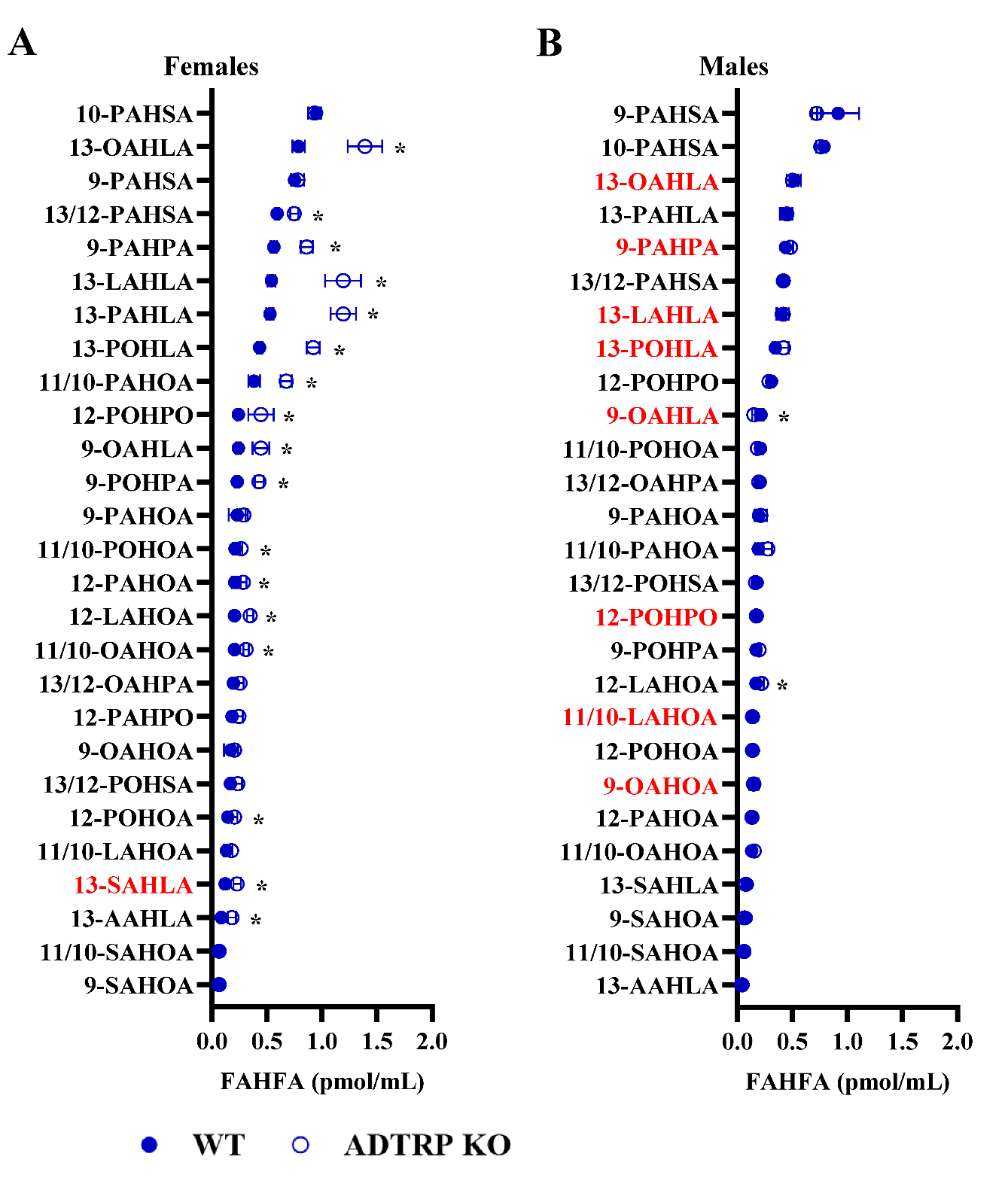
**Figure S1**. Absolute levels of individual FAHFA regioisomers in female (A) and male (B) WT and ADTRP KO CON mice. Data are means ± SE (*n* = 6-7). *Statistically significant vs. WT; *p* ≤ 0.05 (t-test). The FAHFA regioisomers significantly affected by exercise are highlighted in red.

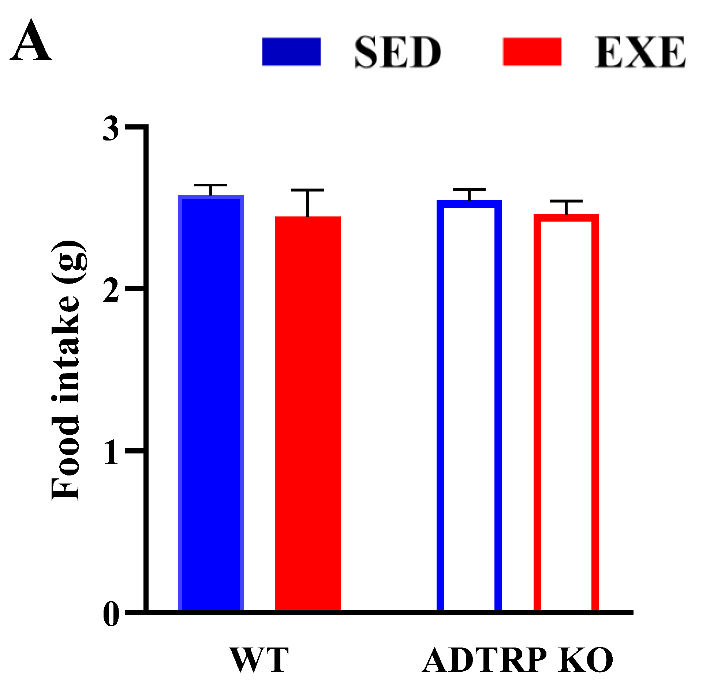

**Figure S2.** Average food intake in SED and EXE mice from the WT and ADTRP KO cohorts during the 7-week experimental period. Data are means ± SE (*n* = 6-7).

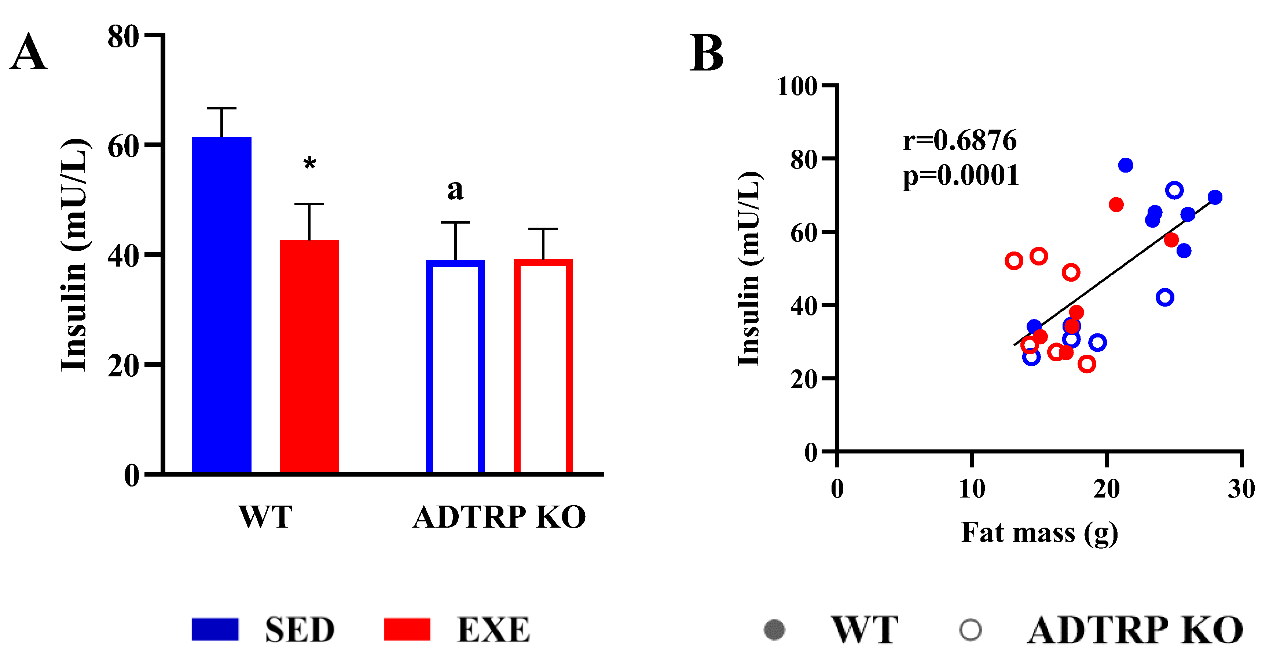

**Figure S3**. Plasma insulin levels in WT and ADTRP KO mice (A). ^a,^*Significant difference vs. WT and SED, respectively; *p* ≤ 0.05 (two-way ANOVA). Data are means ± SE (*n* = 6-7). Relationships between body fat mass and plasma insulin levels (B). The graph shows the Pearson correlation coefficient (r), and the statistical significance of the correlation.

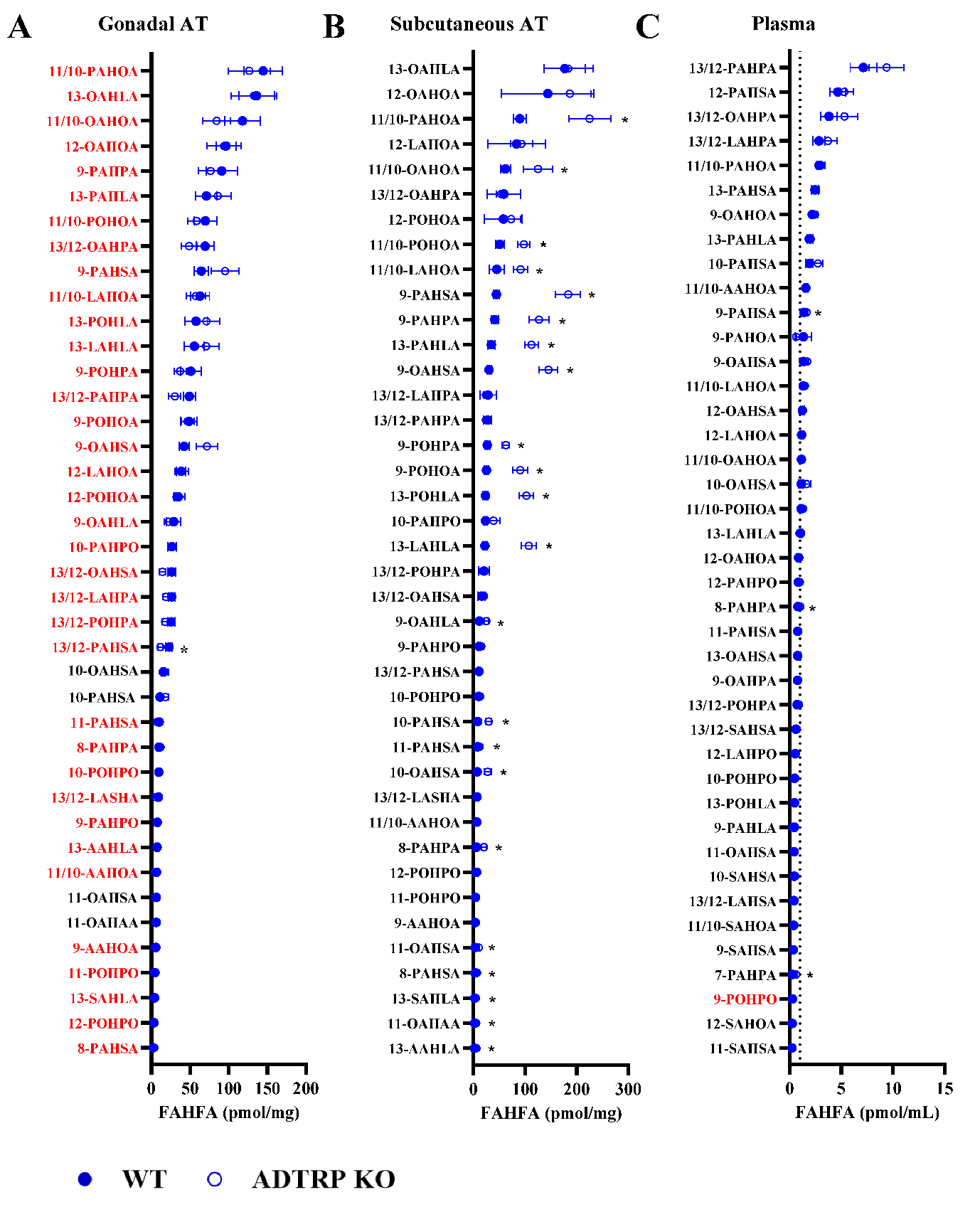
**Figure S4**. Absolute levels of individual FAHFA regioisomers in gonadal AT (A), subcutaneous AT (B), and plasma (C) in WT and ADTRP KO SED mice fed HFD. Data are means ± SE (*n* = 6-7). *Statistically significant vs. WT; *p* ≤ 0.05 (t-test). The FAHFA regioisomers significantly affected by exercise are highlighted in red.

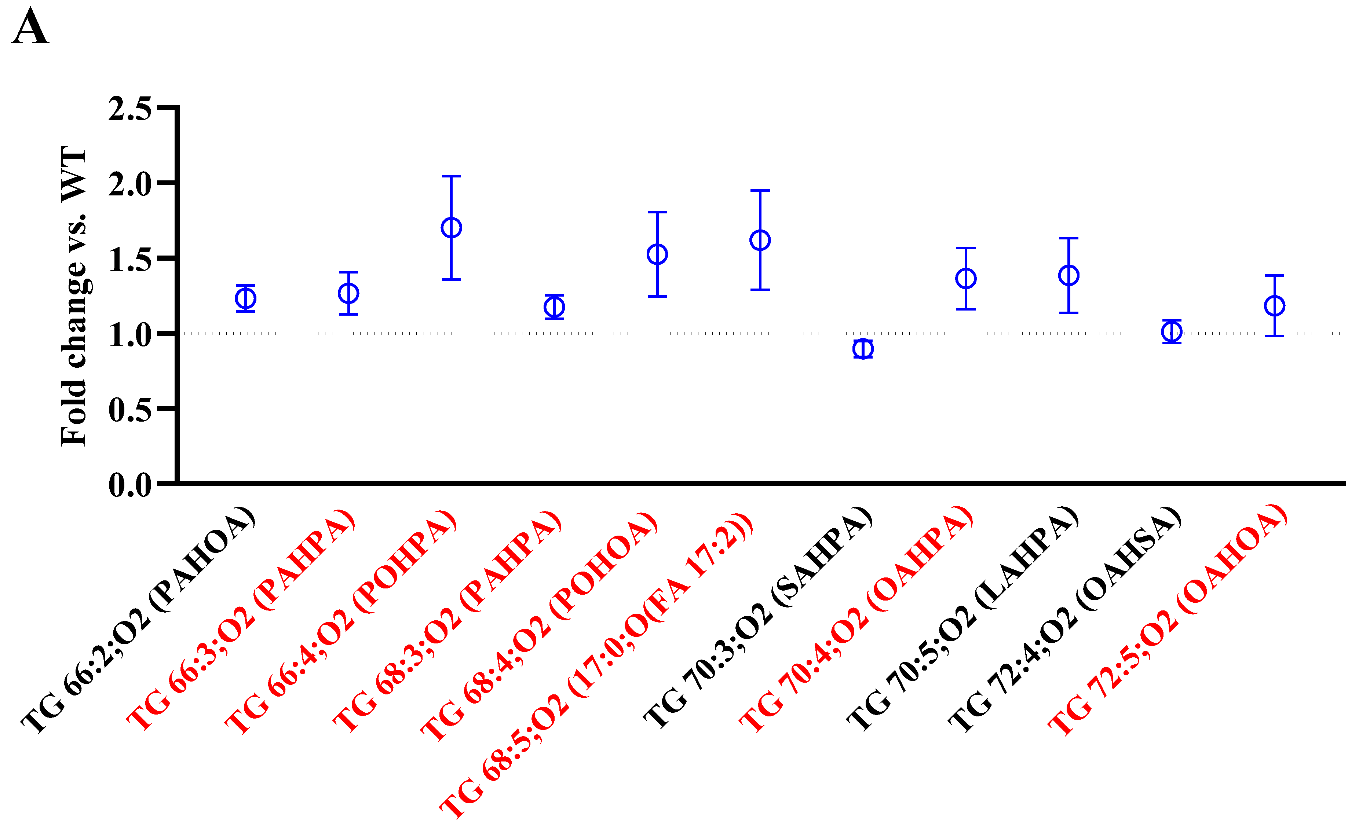

**Figure S5.** The levels of different TG EST in gonadal AT of ADTRP KO SED mice relative to WT counterparts. Data are means ± SE (*n* = 6-7). *Statistically significant vs. WT; *p* ≤ 0.05 (t-test). The individual TG EST on which exercise had a significant effect in WT mice are highlighted in red.
